# Metric-dependent genomic diversity reveals phylogenetic and ecological structure in saproxylic beetles

**DOI:** 10.64898/2026.08.30.748084

**Authors:** Rama Sarvani Krovi, Nermeen R. Amer, Alicja Wierzbicka, Radosław Plewa, Marcin Kadej, Tomasz Jaworski, Adrian Smolis, Tomasz Szmatoła, Maria Oczkowicz, Łukasz Kajtoch

**Affiliations:** Institute of Systematics and Evolution of Animals, Polish Academy of Sciences, Kraków, Poland; Entomology Department, Faculty of Science, Cairo University, Giza, Egypt; Department of Animal Molecular Biology, National Research Institute of Animal Production, Balice, Kraków, Poland; Forest Research Institute, Department of Forest Protection, Sękocin Stary, Raszyn, Poland; University of Wrocław, Department of Invertebrate Biology, Evolution and Conservation, Wrocław, Poland; Department of Basic Science, University of Agriculture in Krakow, Kraków, Poland

**Author notes:** joint first authors (Rama Sarvani Krovi, Nermeen R. Amer). Corresponding author:* Rama Sarvani Krovi, (48 12) 422-70-06, ul. Sławkowska 17 31-016, Kraków.

**Keywords:** comparative population genomics, ddRAD-seq, genetic diversity, host-tree association, macrogenetics, phylogenetic signal, saproxylic beetles

## Abstract

1. Comparative studies can reach different conclusions when genomic-diversity metrics use different denominators.
2. We analysed standardized ddRAD-seq data from 759 individuals, 149 species-by-site population records and 26 saproxylic beetle species across eight Polish forest complexes.
3. Species identity and sampling hierarchy explained most population-level variation. Deciduous-associated species had higher variant-site expected heterozygosity than conifer-associated species.
4. All-position nucleotide diversity showed stable phylogenetic signal, whereas occurrence tier and exploratory forest-management and protection contrasts showed no consistent associations.
5. Genomic-diversity metrics are not interchangeable; comparative assessments should report their denominator and test lineage sensitivity.

## Introduction

Intraspecific genetic diversity contributes to population persistence and adaptive potential, yet it remains the least understood level of biodiversity. Macrogenetics seeks general explanations for variation in genetic diversity by comparing many populations and species across space (Blanchet et al. 2017; Leigh et al. 2021). Candidate explanations include abundance, geographic range, ecological specialization, dispersal, evolutionary history and anthropogenic disturbance. Their relative importance remains uncertain because comparative datasets often combine different markers, sampling designs and analytical pipelines.

Genome-wide data generated under a common workflow can reduce some of this heterogeneity (Yahara et al. 2010, Coates et al. 2018). They do not, however, eliminate differences among diversity metrics. Nucleotide diversity calculated across all callable positions incorporates invariant sites and represents per-site sequence diversity, whereas heterozygosity reported only among retained variant sites is conditional on discovery and filtering (Schmidt et al. 2021). Other metrics are mostly derived from one of these two basic variables (i.e. inbreeding (F_IS_) is measured based on observed and expected heterozygosity), and are not independent measures of diversity. Treating these responses as interchangeable can obscure whether a predictor is associated with a general genomic property or with a particular metric and denominator (Brault et al. 2026).

Saproxylic beetles provide a useful assemblage for testing ecological and evolutionary hypotheses. They depend on deadwood and tree-related microhabitats, contribute substantially to decomposition, and include both abundant taxa (some treated as pests) and rare specialists of conservation concern (Stokland et al. 2012; Seibold et al. 2015). Host-tree association, microclimatic niche, and species distribution (range) may influence effective population size and population connectivity, which can be estimated using genomic data. At the same time, mutation rate, life-history traits and demographic history may be conserved among related lineages, generating similar genetic patterns among congeners or confamilial species.

Forest management may also be one of the factors that correlate with beetle genetic diversity because it alters the quantity, quality and continuity of deadwood, although human-made alterations are relatively recent on an evolutionary timescale and therefore their effect may not yet be detectable.

Previous studies have already demonstrated that both the geographic location and management regime of forests influence the genetic diversity of saproxylic beetles (Amer et al. 2026), and that deadwood diversity, together with the duration of forest protection, are key determinants of genetic diversity in this ecological group (Krovi et al., 2026). However, these studies did not examine how phylogenetic affinity, biogeographic history, microclimatic preferences, trophic specialization, and other ecological traits shape patterns of genetic diversity across multiple saproxylic beetle species. The novel contribution of the present study is therefore a cross-taxonomic, assemblage-scale assessment of the effects of species traits, phylogenetic affiliation, and forest management on genome-wide within-population genetic diversity across 26 saproxylic beetle species. This addresses a macrogenetic question that has not been explored in previous studies. Specifically, we tested the following three hypotheses:

H1. Species with high population abundance are expected to exhibit greater genomic diversity than habitat specialists, particularly taxa associated with old-growth forest remnants, owing to greater habitat connectivity and larger, more persistent populations.

H2. Species associated with deciduous deadwood are expected to exhibit greater genomic diversity than conifer specialists, reflecting the historical predominance of broadleaved forests and contrasting outbreak dynamics.

H3. Shared evolutionary history is predicted to structure genomic diversity, most clearly for all-position nucleotide diversity, whereas variant-site heterozygosity may be more responsive to recent ecological and demographic processes.

## Materials and Methods

### Study system and sampling

We analysed 759 beetle specimens representing 26 species and 14 families Table S1. Samples formed 149 species-by-site population records across 37 sites in eight Polish forest complexes: Augustów (AF), Białowieża (BF), Carpathian Mountains (CM), Dolina Baryczy (DB), Holy Cross Mountains (HM), Knyszyń (KF), Oder Forests (OF) and Silesian Forests (SF). Three complexes were classified as primeval (BF, CM, HM) and the remaining five as managed, shown in Figure 1. Because every site within a complex had the same management category, management was completely nested within the forest complex. We consequently treat management as an exploratory between-forest comparison. Population records contained five individuals on average; species were represented by 3–10 population records.

**Figure 1.**
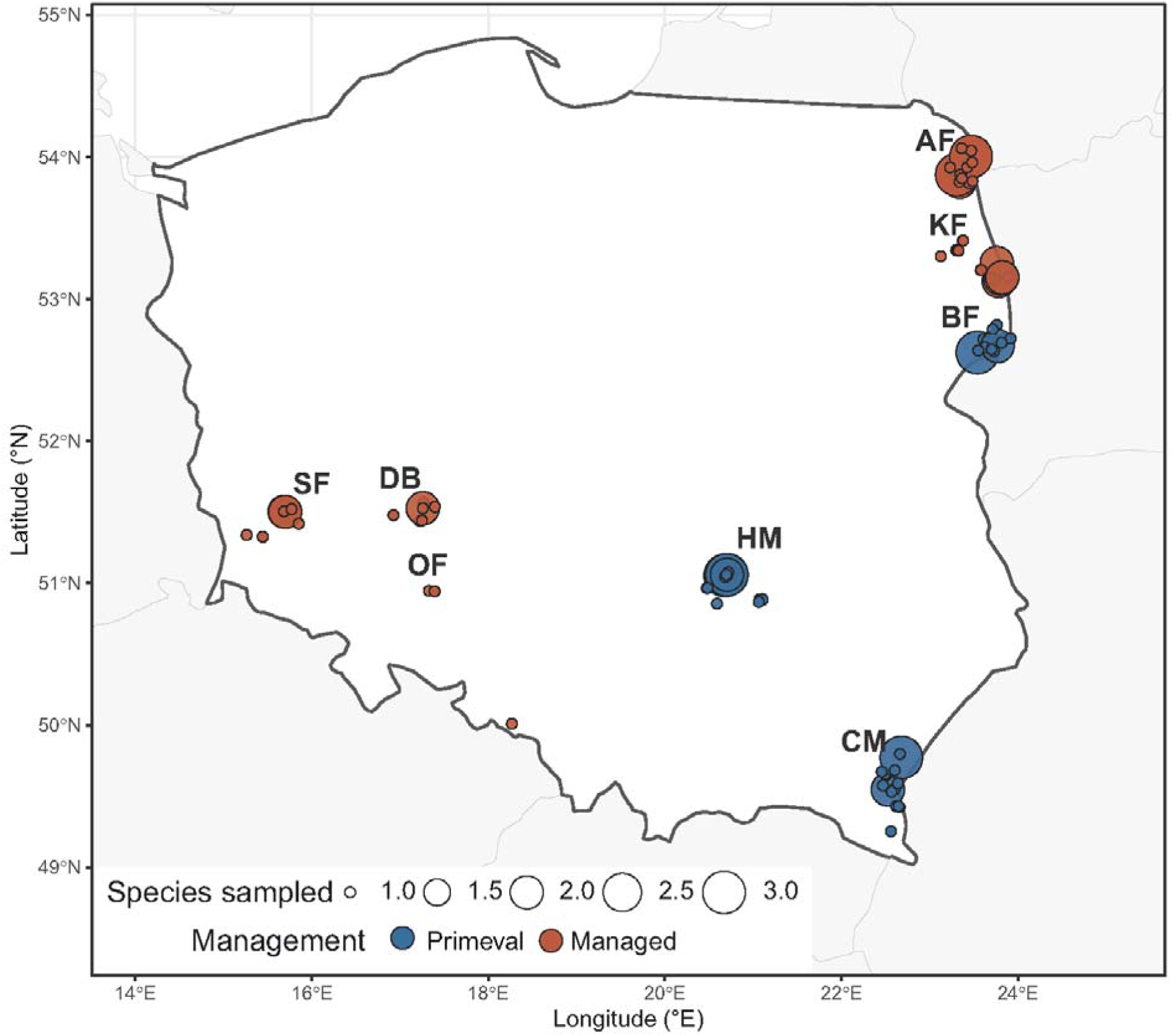
Distribution of sampled beetle populations across Poland. Points represent 105 distinct recorded coordinates among 149 species-population records. Point size indicates the number of population records sharing the same recorded coordinates, and colour indicates the management category of the forest complex.

### Traits data collection

Species traits were compiled from the project trait database (organised based on literature, e.g. Seibold et al. 2015, Wende et al. 2017, Hagge et al. 2021, including expert knowledge of co-authors of this study), and checked for consistency within species. Candidate traits included: “family” (taxonomic assignment to the beetle family), “abundance” (having populations with numerous individuals vs rare), “distribution” (boreal vs temperate vs boreal & temperate), “conservation protection status” (protected vs not), “shade tolerance” (moist and cold vs dry and warm), “economic value” (neutral vs pest, in respect to timber production), “trophic guild” (invertivorous vs xylovorous vs fungivorous), “microhabitat” (living under bark vs wood vs tree cavities vs fungi), “settlement” (deadwood decay: fresh vs old), “tree type” (coniferous vs deciduous vs various =both), “host tree” (host tree species), “body size” (imago: small <10 mm, medium 10-20 mm, large >20 mm), “dispersal ability” (low vs moderate vs high), “voltinism” (uni-vs multivoltine= number of generations/year). Categorical predictors were retained for comparative testing only when every level contained at least five species. This rule retained the “abundance”, “host-tree” category and “microclimatic” niche among the hypothesis-level predictors. The complete trait list and level counts are provided in Supporting Tables S2–S3.

### DNA isolation, ddRAD sequencing and variant processing

High molecular weight DNA was extracted using the Sherlock AX kit (A&A Biotechnology). ddRAD-seq libraries were prepared following Bayona-Vásquez et al., (2019) with modifications by Amer et al., (2026). Double-digest RAD libraries were prepared using NheI and EcoRI, size-selected to 400–600 bp and sequenced as 150-bp paired-end reads on an Illumina NovaSeq X Plus. Reads were demultiplexed and quality filtered before species-specific de novo assembly in Stacks v2.68 (Rochette et al. 2019). The documented workflow used a minimum stack depth of three and species-specific assembly optimization. An initial Stacks populations run required loci in at least two populations and at least 80% of individuals per population. PLINK filtering (Purcell et al. 2007) then retained markers with minor-allele frequency ≥0.03 and no more than 30% missing genotypes; individuals with >50% missing data were screened before the final whitelist-based populations run.

### Genetic diversity metrics

Population statistics were extracted from the Stacks populations.sumstats_summary.tsv files. The primary response, nucleotide diversity (π), was taken from the “All positions” section and therefore included invariant callable sites in the denominator. Observed heterozygosity (H□), expected heterozygosity (H□) and F_IS_ were taken from the “Variant positions” section and describe retained variable sites. We retained both all-position and variant-position values in the audit dataset, but did not treat the denominators as equivalent. Allelic richness was excluded because the available values exceeded the theoretical maximum for correctly rarefied biallelic SNPs. Per-species mean callable-site counts and sample coverage, together with sampling design and species-mean diversity, are reported in Supporting Table S2.

### Phylogeny

The 26-species phylogeny was derived from the Open Tree of Life using rotl (Michonneau et al. 2016), then processed using ape (Paradis & Schliep 2019) and phytools (Revell 2012) after taxonomic reconciliation. Polytomies were resolved deterministically and Grafen branch lengths were applied where the induced tree was not ultrametric (Grafen 1989). Because this synthetic phylogeny is not time calibrated, phylogenetic estimates are interpreted as comparative structure rather than evolutionary rates. Pagel’s λ and Blomberg’s K were calculated for raw and log-transformed metrics where transformation was defined, using 1,000 permutations for K. Sensitivity analyses were repeated after excluding Rhysodes sulcatus, a relict species associated with European primeval forests (Kostanjsek et al. 2018), the sole representative of the suborder Adephaga (all other taxa belonging to Polyphaga; McKenna et al. 2019), which exhibited substantial phylogenetic leverage.

### Statistical analysis

Analyses were conducted at both population and species levels in R version 4.3.0 (R Core Team 2023)

1. Population-level Linear Mixed Models (LMMs): LMMs were fitted using lme4 (Bates et al. 2015) to test ecological predictors while controlling for hierarchical structure via nested random effects (Family/Species and Forest/Site) (Bolker et al., 2009). The candidate model set was restricted a priori to: (i) Intercept-only null, (ii) Occurrence tier, (iii) Host-tree category, (iv) Microclimatic niche, (v) Standardized body size, and (vi) Occurrence + Host-tree category. Models were fitted via Maximum Likelihood (ML) and ranked using corrected Akaike Information Criterion (AICc) with MuMIn package (Bartoń 2010; Burnham & Anderson 2004). To avoid over-fitting species-level traits evaluated on sub-sampled populations, AICc small-sample corrections were calculated using the number of species (N = 26) rather than total population records (N = 149). The top-performing models were re-fitted using Restricted Maximum Likelihood (REML). To facilitate convergence for small numerical values (e.g., π), responses were scaled by a constant prior to modeling, and parameter estimates were back-transformed to original units for reporting. Fixed-effect degrees of freedom and P-values were obtained using Satterthwaite’s approximation as implemented in lmerTest (Kuznetsova et al. 2017). Marginal and conditional (R²) values were calculated using performance (Lüdecke et al. 2021).
2. Species-level Phylogenetic Generalized Least Squares (PGLS): Species-level metric means were evaluated using PGLS models in caper (Orme et al. 2011), estimating Pagel’s λ by Maximum Likelihood. Response variables π, H□, and H□ were log-transformed; F_IS_ was analyzed on its original scale due to zero/negative values. Single-predictor models evaluated occurrence tier, host-tree category, microclimatic niche, and body size. Benjamini–Hochberg (BH) false discovery rate corrections were applied to p-values across single-predictor models within each response. Sensitivity runs evaluated the full 26-species dataset against the 25-species subset excluding *R. sulcatus*.
3. Forest Management Evaluation: Forest management category was excluded from candidate ecological model selection to prevent confounding effects. Instead, management was evaluated as an exploratory fixed factor (primeval vs. managed) within an LMM retaining the Forest/Site random hierarchy. An additional exploratory LMM evaluated protected vs. unprotected population records (74 vs. 75 populations). Model diagnostics included residual normality checks, variance inflation factors (VIF), spatial autocorrelation tests (Moran’s I), and leave-one-species-out cross-validation.

## Results

### Population-Level Baseline and Multi-Level Variance (Null Models)

In population-level linear mixed-effects models (LMMs), the intercept-only null model ranked first across all four genomic diversity metrics. For all-position nucleotide diversity (□), the second-ranked candidate model was standardized body size (ΔAICc = 2.21), whereas the occurrence tier and host-tree category yielded ΔAICc values of 3.78 and 4.62, respectively. For variant-site observed (H□) and expected (H□) heterozygosity, host-tree association ranked second but remained behind the null (ΔAICc =1.30 for H_o_; ΔAICc = 1.91 for H_e_). The null model was also preferred for the inbreeding coefficient (F_IS_). Conditional R² values for null models were high (0.69-0.92), showing that species identity and multi-level site clustering account for most variance across all metrics rather than ecological predictor effects.

### Occurrence Tier and Macroecological Prevalence (H1)

Occurrence pattern did not consistently predict any genomic metric after false discovery rate (BH) correction (Table 1). For all-position π, common species showed an initial positive unadjusted contrast relative to rare species (β= 0.267, raw P = 0.034), but this effect was not retained following Benjamini–Hochberg correction (adjusted P = 0.172). For variant-site H□ and H□, common and pest species tended toward lower point estimates than rare species, though no contrasts reached statistical significance. F_IS_ contrasts across occurrence tiers were consistently weak and non-significant.

**Table 1.** Principal species-level ecological-trait results.

| Predictor | Response | Estimate | Raw P | BH P | Sensitivity | Conclusion |
| --- | --- | --- | --- | --- | --- | --- |
| Host: deciduous vs coniferous | H<sub>□</sub> (log) | $\beta = 0.212$; SE = 0.066 | 0.0041 | 0.036 | $\beta = 0.216$; P = 0.0027 without *R. sulcatus* | Supported |
| Host: deciduous vs coniferous | H<sub>□</sub> (log) | $\beta = 0.286$; SE = 0.109 | 0.015 | 0.103 | Positive but not retained after correction | Suggestive |
| Host: deciduous vs coniferous | $\pi$ (log) | $\beta = -0.056$; SE = 0.116 | 0.635 | 0.635 | No relationship | Unsupported |
| Occurrence: common vs rare | $\pi$ (log) | $\beta = 0.267$ | 0.034 | 0.172 | Not retained after correction | Suggestive |
| Occurrence tier | H<sub>□</sub>, H<sub>□</sub> and F<sub>IS</sub> | Directions inconsistent | > 0.05 | > 0.05 | No corrected association | Unsupported |

Note: $\pi$ was calculated across all callable positions. H<sub>□</sub> and H<sub>□</sub> were calculated among retained variant positions. BH correction was applied within response.

### Interspecific Trait Specialization and Host-Tree Association (H2)

In species-level PGLS models, host-tree association significantly predicted variant-site H□ (Table 1; Figure 2). Deciduous-associated species exhibited significantly higher H□ than conifer-associated species (log-scale β = 0.212, SE = 0.066, P = 0.0041, adjusted P = 0.036). In contrast, host-tree association showed no detectable effect on all-position □ (β = −0.056, SE = 0.116, P = 0.635). Although host-tree category and microclimatic niche were partially correlated, generalized variance inflation factors (GVIF 1.21 and 1.59) confirmed predictor separability. Microclimatic niche alone did not predict any metric (Hₑ β = 0.067, P = 0.444; Hₒ β = 0.111, P = 0.438; π β = 0.080, P = 0.496), and species utilizing “various” host trees exhibited expected heterozygosity levels comparable to conifer specialists, confirming that elevated Hₑ specifically tracks deciduous host-tree association independently of shade or moisture preferences.

**Figure 2.**
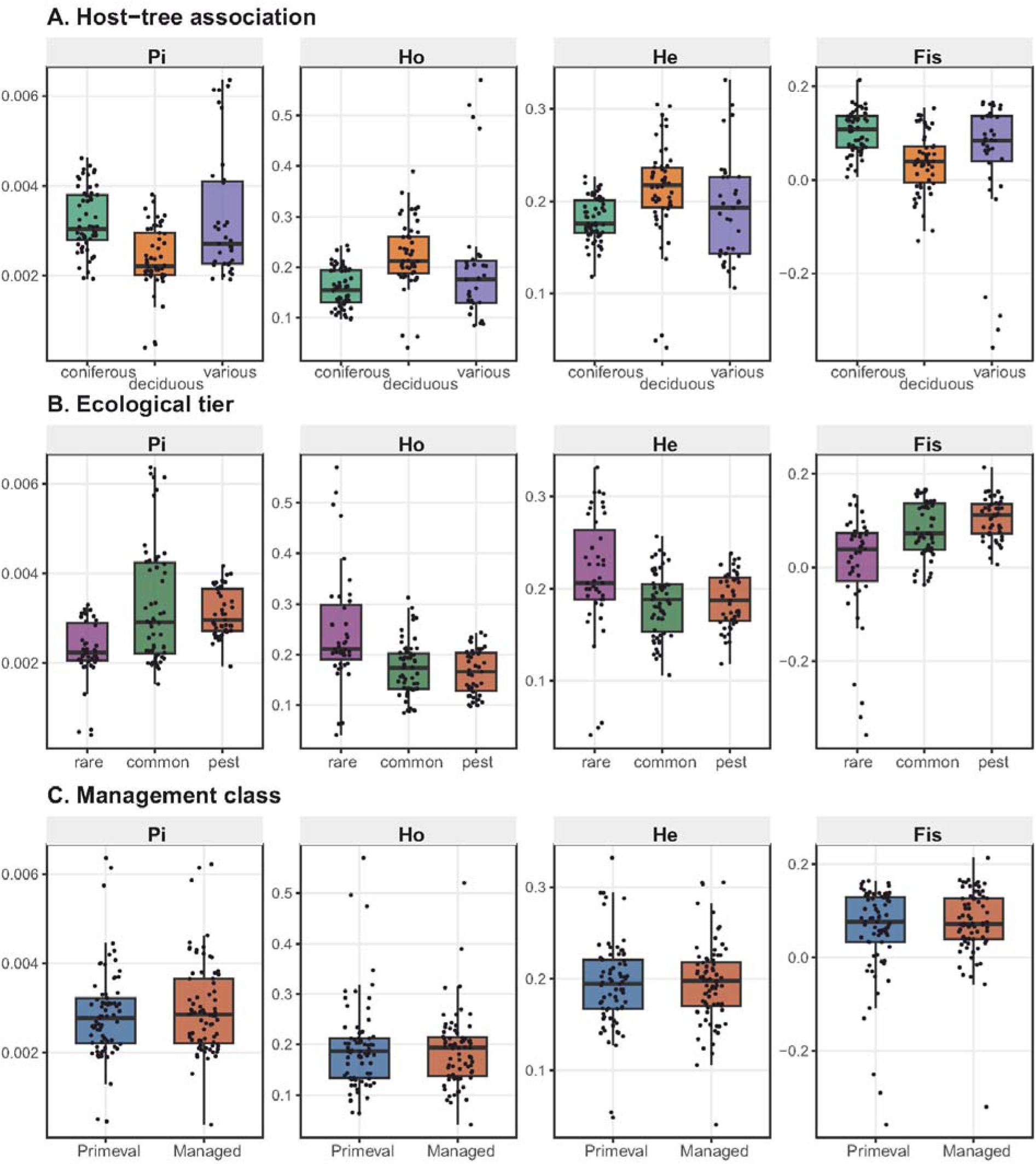
Genetic diversity parameters (π, Ho, He, F_IS_) grouped by (A) host-tree preference, (B) occurrence tier, and (C) forest management regime. Box plots display medians and interquartile ranges, with individual species observations superimposed.

### Evolutionary History and Metric-Specific Phylogenetic Signal (H3)

All-position □ exhibited strong, consistent phylogenetic signal across both log and raw scales (Table 2; Figure 3). On the log scale, Pagel’s λ was 0.583 (P = 0.0099) for the full 26-species assemblage and remained stable at λ = 0.580 (P = 0.0126) in sensitivity tests excluding *Rhysodes sulcatus*. Raw-scale estimates were similarly stable (λ = 0.472, P = 0.032 for full data; λ = 0.468, P = 0.037 without *R. sulcatus*). This strong signal in all-position π contrasted sharply with variant-site metrics, which showed weak or lineage-sensitive signals. Variant-site H□ displayed a lineage-sensitive signal in the full dataset (λ = 0.657, P = 0.0296), but this signal was not retained after removal of *R. sulcatus* (λ = 0.257, P = 0.232). Variant-site H_e_ did not exhibit significant Pagel’s λ in either dataset, and its Blomberg’s K signal was non-significant in the sensitivity run.

**Table 2.**
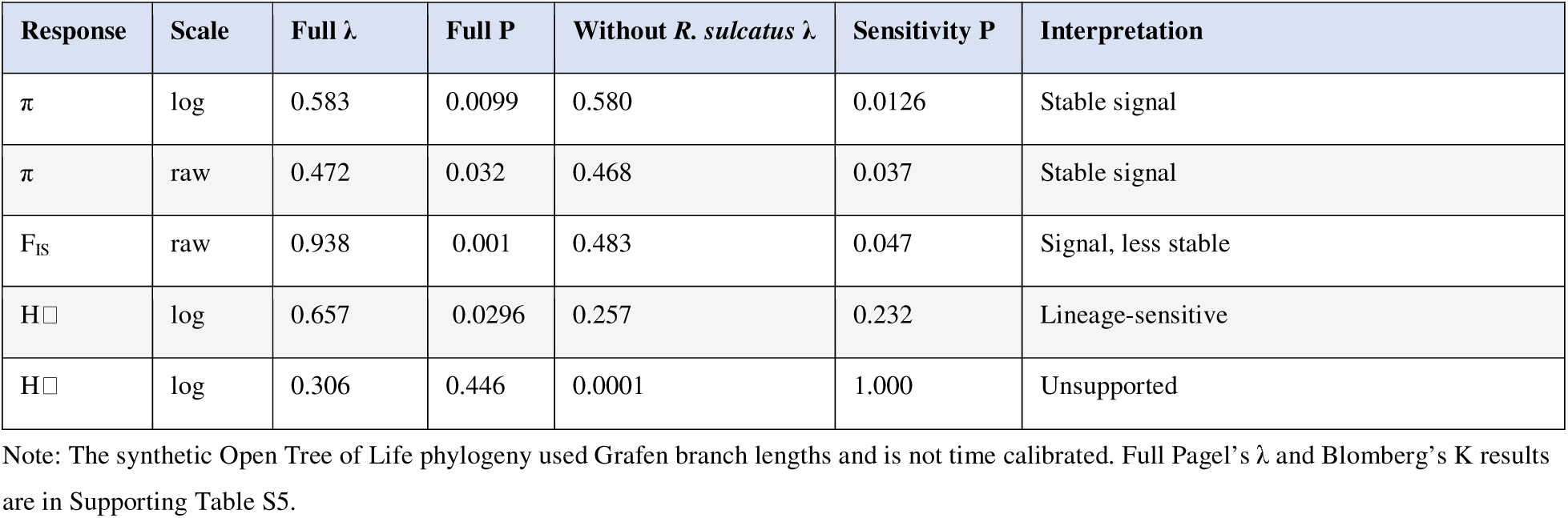
Summary of phylogenetic-signal tests and lineage-removal sensitivity.

**Figure 3.**
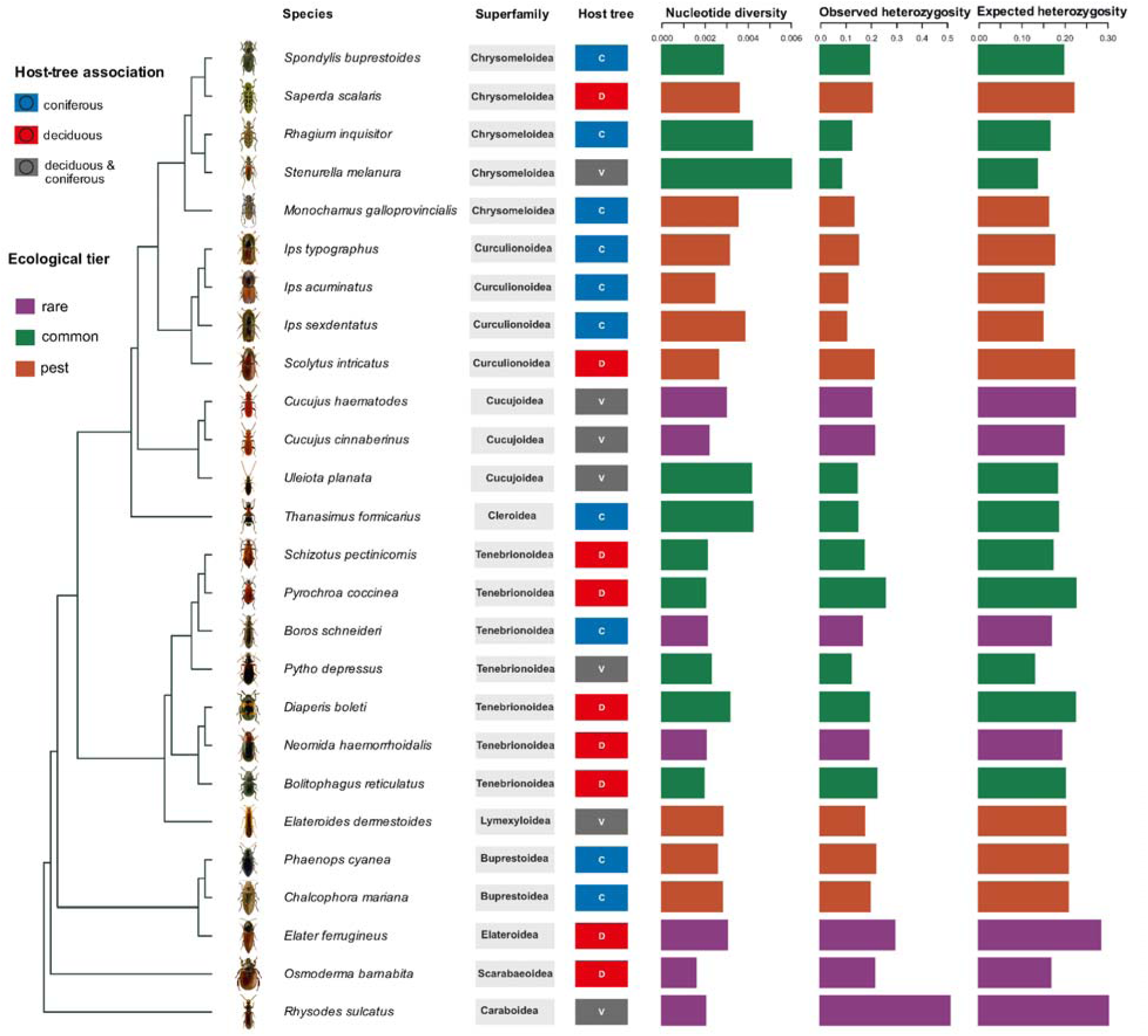
Phylogeny and species-mean genomic diversity for 26 saproxylic beetles. Bars show all-position nucleotide diversity (π) and variant-site observed and expected heterozygosity (H□ and H□). Colours indicate occurrence tier.

### Forest Management and Landscape Nesting

In exploratory LMMs, managed forest complexes showed slightly higher point estimates across all four metrics, but all 95% confidence intervals overlapped zero (0.00011 for π, 95% CI: −0.00008 to 0.00030; 0.005 for Hₒ, 95% CI: (−0.014 to 0.025; 0.010 for Hₑ, 95% CI: −0.007 to 0.027; 0.009 for F_IS_, 95% CI: −0.003 to 0.020). In exploratory models comparing protected versus unprotected population records (74 vs. 75 records), protected status showed no detectable association with any metric (H□ β = 0.005, 95% CI −0.009 to 0.018, P = 0.497; H□ β = −0.001, 95% CI −0.011 to 0.009, P = 0.803; F_IS_ β = −0.011, 95% CI −0.023 to 0.000, P = 0.058). Because the management category was represented by eight forest complexes and was completely nested within forest complex identity, these LMM estimates might reflect broad spatial variation among sampled landscapes rather than direct management or protection effects.

## Discussion

Previous research on saproxylic beetles examined associations between ecological and morphological traits and extinction risk, community composition and environmental responses (Seibold et al. 2015; Hagge et al. 2021), without testing how these traits relate to within-species genomic diversity. Our study links these approaches by comparing multiple measures of genomic diversity across 26 species spanning several beetle families and ecological strategies. Although limited species coverage constrains broad generalisation across Coleoptera, it provides a comparative test of ecological and phylogenetic associations with population genetic diversity in saproxylic beetles.

Our comparative analysis reveals that genomic diversity is shaped by distinct, scale-dependent processes. Variance in genomic diversity is primarily structured by baseline species identity, with ecological traits explaining specific, metric-dependent fractions of diversity. Crucially, our findings show that occurrence patterns do not consistently dictate genomic diversity (H_1_), whereas host-tree specialization specifically drives variant-site expected heterozygosity (H_e_) in broadleaf-associated taxa (H_2_). Furthermore, deep evolutionary history strongly constrains all-position nucleotide diversity (π), whereas variant-site heterozygosity reflects recent ecological specialization (H_3_).

### Occurrence patterns do not dictate genomic diversity (H_1_)

Contrary to the hypothesis that species with populations comprising numerous individuals maintain elevated genomic diversity relative to rare or localized specialists, abundance tier showed no significant association with any metric after false discovery rate correction. While abundant species exhibited a non-significant positive trend for all-position π, variant-site heterozygosity (H_o_ and H_e_) tended to be slightly lower in abundant (including pest) species than in rare taxa. This disconnect highlights that macroecological prevalence (geographic range and local abundance) does not directly reflect evolutionary effective population size (Ne). Specialized, habitat-restricted saproxylic beetles may maintain historically stable, highly connected populations in primeval forest refugia, buffering them against rapid genetic drift despite localized distribution ranges (Seibold et al. 2015). Conversely, abundant (including pest) species frequently undergo local population outbreaks followed by bottlenecks- (Lantschner & Corley 2023), inducing recurring founder effects that temporarily inflate variant-site homozygosity, and may have implications for the genomics of irruptive beetles (Mykhailenko et al. 2024). Finally, it should be emphasized that not all rare species have restricted distributions and, conversely, some range-restricted species may have abundant populations. Therefore, assigning taxa to two simple categories can be ambiguous, and abundance should not be treated as equivalent to commonness or widespread distribution. More quantitative measures of population abundance and range occupancy would provide a stronger test of these relationships.

### Host-tree association drives variant-site expected heterozygosity (H2)

Species associated with deciduous host trees exhibited significantly higher expected heterozygosity than conifer specialists. This relationship was independent of microclimatic niche preferences (shade and moisture), as confirmed by low variance inflation factors. Temperate forest habitats in Central Europe represent ancestral, structurally complex broadleaf ecosystems that have maintained continuous long-term microhabitat availability throughout post-glacial recolonization (Roberts et al. 2018). Conifers were also present in these forests, although the dominance of pines and spruces is relatively recent and was caused by human forest management (Jansen et al. 2017). The broad dietary and structural niches provided by decaying broadleaf wood support larger, long-term effective population sizes in deciduous specialists (Seibold et al. 2015; Ulyshen 2018), which is why most primaeval relics are among taxa associated with deciduous trees. In contrast, conifer-associated saproxylic beetles historically occupied restricted boreo-montane refugia prior to modern commercial forestry expansions (Kajtoch et al. 2022, Krovi et al. 2025). The lower H_e_ observed in conifer specialists likely reflects historical demographic bottlenecks rather than modern forestry practices, as recent conifer plantation expansions have occurred too recently to rebuild standing variant-site heterozygosity (Lachat et al. 2012).

### Evolutionary history and the denominator distinction (H3)

Previous beetle phylogenomic studies have examined deep evolutionary relationships and diversification across Coleoptera (Zhang et al. 2018; McKenna et al. 2019), while studies within particular herbivorous groups have linked diversification to host-plant use (McKenna et al. 2009). Until now, however, no research has linked phylogeny with intraspecific genomic diversity across multiple taxa for deadwood-dwelling organisms.

Our results strongly support H_3_ for sequence-level divergence while demonstrating that phylogenetic signal is metric-dependent. All-position nucleotide diversity retained a strong, stable phylogenetic signal, remaining invariant to the removal of the basal lineage *Rhysodes sulcatus*. In contrast, variant-site heterozygosity (H_e_) showed no significant phylogenetic signal. This distinction points at a methodological denominator distinction in restriction site-associated DNA sequencing (RAD-seq) (Harvey et al. 2016). All-position π incorporates invariant callable sites into the denominator, causing it to reflect deep, genome-wide mutation accumulation across evolutionary time (Nei & Li 1979; Lynch & Conery 2003). Because mutation accumulation is constrained by lineage age and historical metabolic rates, π exhibits strong phylogenetic conservatism (Pagel 1999; Blomberg et al. 2003). Conversely, variant-site metrics (H_o_ and H_e_) restrict the denominator to retained polymorphic SNPs, isolating active allele frequency distributions that respond rapidly to recent ecological selection, host specialization, and population dynamics (Luikart et al. 2003; Allendorf et al. 2012). Inbreeding (F_IS_) also retained evidence of signal after lineage removal, but it is a composite of H□ and H□ and should not be interpreted as an independent measure of genetic diversity. Evaluating both nucleotide diversity and heterozygosity simultaneously prevents erroneous conclusions regarding species-wide genetic vulnerability.

### Spatial nesting masks direct forest management effects

Exploratory comparisons between managed and primeval forest complexes revealed no consistent management effects across any diversity metric. Although managed forests showed slightly higher point estimates, the 95% confidence intervals overlapped zero in all models. This absence of a direct management signal stems from spatial nesting: because management status was uniform within entire forest complexes, management effects were completely confounded with broad-scale geographic identity and regional forest history (Dormann et al. 2007). Population-level linear mixed models confirmed that random site and species identity accounted for 69–92% of total variance.

First, although our sampling encompassed both primeval and managed forests, populations of rare species and most common forest specialists in commercially managed forests were almost exclusively found in relatively well-preserved old-growth stands, including not only nature reserves but also forest patches that had remained free from logging for extended periods (Amer et al. 2026). In contrast, only abundant species, including pests, were consistently sampled in intensively managed stands. Second, forests in Central Europe, including those in Poland, comprise a heterogeneous mosaic of stand types, and even commercially managed forests often contain remnants of old-growth habitat (only some being primaeval) (Banaś et al. 2014, Bujoczek et al. 2021). The absence of a detectable management effect may partly reflect this complex forest structure, as well as the necessarily simplified classification of beetle populations into either primeval or managed forest categories. Consequently, evaluating forest management impacts requires finer-scale microhabitat measurements such as local deadwood volume and diversity, or the availability of old trees, rather than categorical landscape designations (Müller et al. 2010).

## Conclusions

The main methodological outcome of this study is that some hypotheses can be evaluated with particular genomic diversity metrics, whereas others remain inconclusive. This is especially important in cross-species de novo SNP datasets, where catalogue size, locus recovery and the pool of retained variants differ among lineages.

Our findings caution against using macroecological rarity or categorical protection status as sole proxies for genomic vulnerability in the conservation of saproxylic beetles (Calix et al. 2018; Eckelt et al. 2018). Rare saproxylic specialists inhabiting broadleaf forest remnants often harbor substantial variant-site heterozygosity, underscoring the conservation value of primeval broadleaf forest fragments (Krovi et al. 2025; Amer et al. 2026). For conservation (rare taxa) or management (pests), the data support lineage-aware monitoring and careful reporting of the genetic component being measured. Maintaining diverse and continuous deadwood resources remains ecologically well motivated (Krovi et al. 2026), but a causal genetic benefit should be tested with direct resource and demographic measurements.

Across the studied saproxylic beetle assemblage, host-tree association and phylogenetic history were linked to distinct patterns of genomic variation. Deciduous-tree association predicted expected heterozygosity among retained variant sites, but not all-position nucleotide diversity. In contrast, π exhibited the strongest and most consistent phylogenetic signal. Abundance patterns did not consistently predict diversity, and the clustered management comparison did not permit a general causal conclusion. These findings emphasize that genomic diversity is not a single property but a hierarchy-dependent characteristic whose interpretation depends on the metric used.

## Acknowledgements and Funding

This research was funded by the National Science Centre, Poland (NCN grant UMO-2021/43/B/NZ9/00991). The authors are sincerely grateful to the Polish nature-protection authorities for granting permission to conduct field work in protected forest complexes and to collect protected species, and to the Polish State Forests National Forest Holding, particularly the Regional Directorates of State Forests in Krosno, Radom, Wrocław, and Białystok, as well as the local foresters.

## Ethical Statement

Collection of protected beetle species and sampling in protected areas was conducted under permissions granted by the Polish Ministry of Environment (DOP-1.61.61.2021.TP 1516071.4994962.3977511 and DOP-1.61.61.2021.TP 1516071.4994586.3977515), the General Directorate of Environmental Protection (DZP-WF.6401.68.2021.AS), and the Regional Directorates of Environmental Protection in Białystok (WPN.6205.74.2021.MM), Rzeszów (WPN.6205.116.2021.MP.2 and WPN.6205.114.2021.ŁL.2), Kielce (WPN.I.6205.1.49.2021.EJP.2) and Wrocław (WPN.6205.143.2021.AR).

## Data Availability Statement

Raw sequencing data and sample metadata are deposited in the NCBI Sequence Read Archive under BioProject PRJNA1399704. Species-specific filtered VCF files are available through RepoOD (https://doi.org/10.18150/HELI6C).

## Author Contributions

Rama Sarvani Krovi: Conceptualization; Investigation; Resources; Formal analysis; Writing – original draft. Nermeen R. Amer: Formal analysis; Investigation; Writing – review & editing. Alicja Wierzbicka and Tomasz Szmatoła: Resources; Writing – review & editing. Radosław Plewa, Marcin Kadej, Tomasz Jaworski and Adrian Smolis: Investigation; Writing – review & editing. Łukasz Kajtoch and Maria Oczkowicz: Conceptualization; Supervision; Funding acquisition; Writing – review & editing.

---

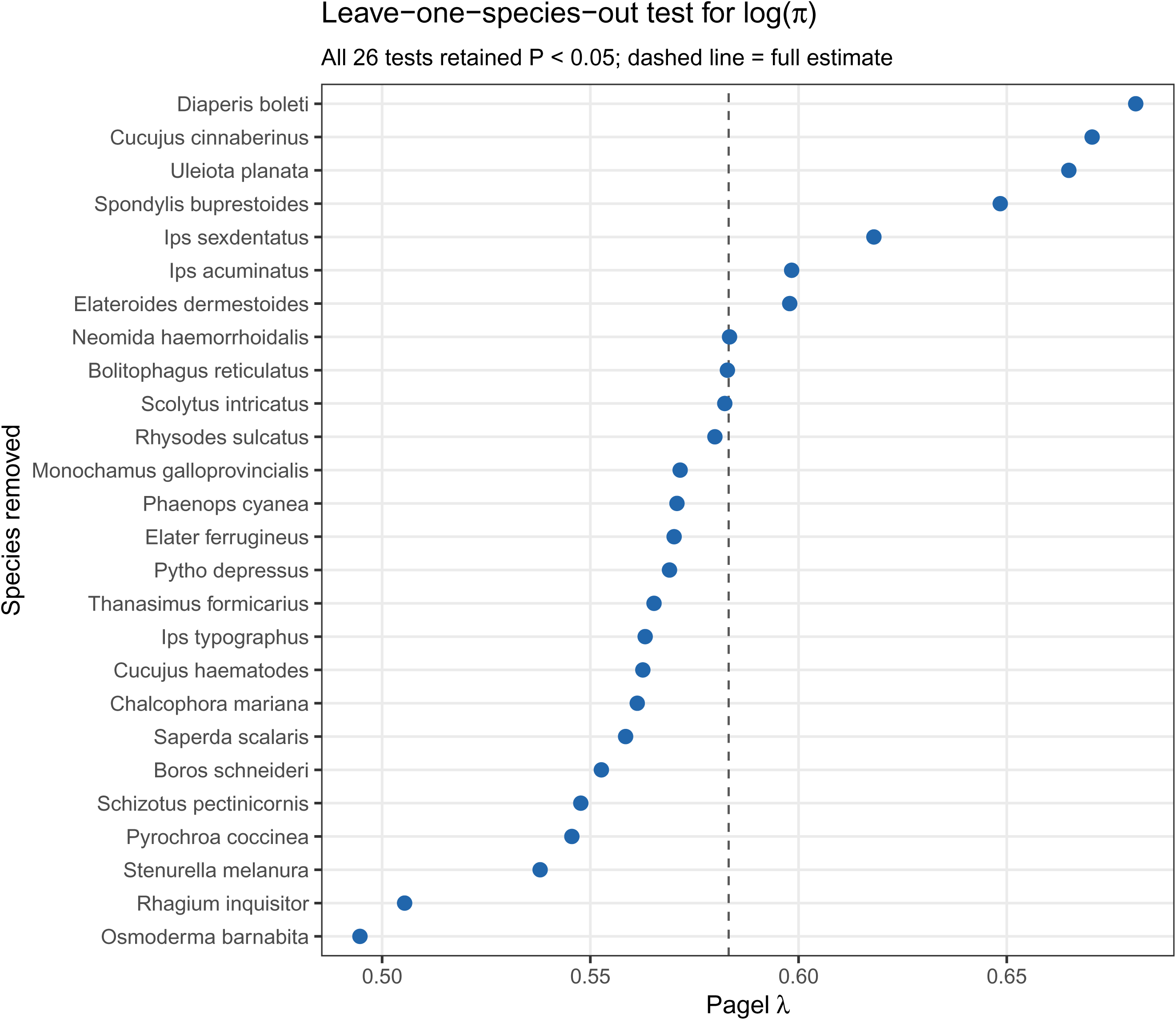

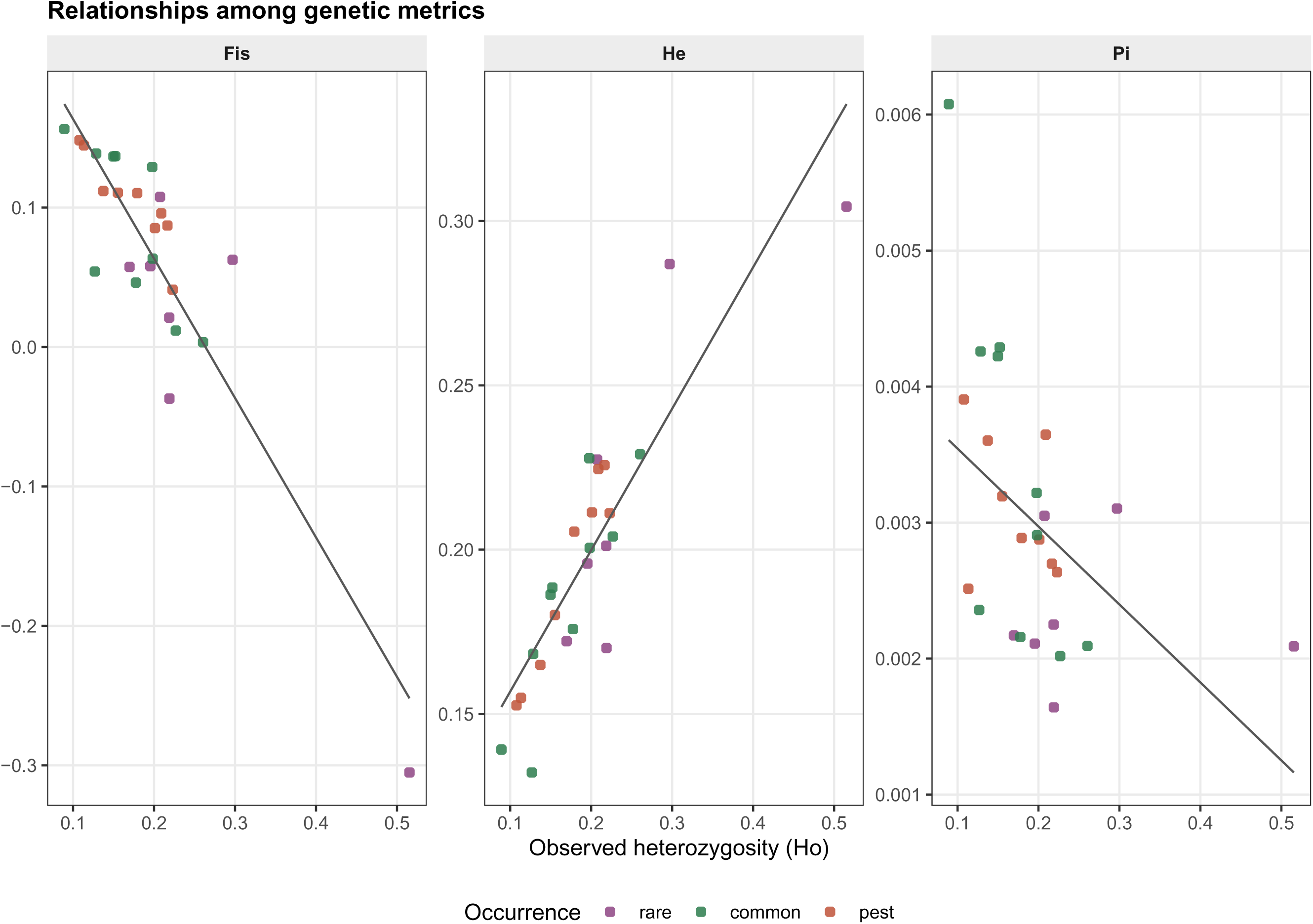

