## Supporting Table S1 for "Metric-dependent genomic diversity reveals phylogenetic and ecological structure in saproxylic beetles"

Supporting Information

Host-tree association and evolutionary history shape components of genomic diversity across a saproxylic beetle assemblage

**Supporting Tables S1–S5**

This document contains the complete supporting tables. Supporting Figures S1–S2 are supplied as separate PDF files.

**Figure S1. Relationships among genomic-diversity metrics.**

**Figure S2. Sensitivity of phylogenetic-signal estimates to removal of Rhysodes sulcatus.**

**Table S1. Population-level analysis dataset, including sampling identifiers, the four analysed genomic metrics, locus-coverage fields and biological predictors.**

| **Species** | **Family** | **Forest** | **Site** | **Management** | **Latitude** | **Longitude** | **Pi** | **Ho** | **He** | **Fis** | **Sites** | **Variant Sites** | **Polymorphic Sites** | **Occurrence** | **Diet** | **Shade tolerance** | **Tree type** | **BL mean** |
| --- | --- | --- | --- | --- | --- | --- | --- | --- | --- | --- | --- | --- | --- | --- | --- | --- | --- | --- |
| Bolitophagus reticulatus | Tenebrionidae | DB | DBd | Managed | 51.4405 | 17.2491 | 0.0015 | 0.1807 | 0.1547 | -0.0159 | 480704 | 4212 | 1744 | common | fungivorous | moist and cold | deciduous | 6.5000 |
| Bolitophagus reticulatus | Tenebrionidae | BF | BFd | Primeval | 52.6334 | 23.7286 | 0.0020 | 0.2119 | 0.2020 | 0.0333 | 485276 | 4241 | 2464 | common | fungivorous | moist and cold | deciduous | 6.5000 |
| Bolitophagus reticulatus | Tenebrionidae | HM | HMc | Primeval | 50.8869 | 21.1055 | 0.0022 | 0.2724 | 0.2228 | -0.0299 | 481948 | 4240 | 2544 | common | fungivorous | moist and cold | deciduous | 6.5000 |
| Bolitophagus reticulatus | Tenebrionidae | CM | CMa | Primeval | 49.6562 | 22.4945 | 0.0020 | 0.2126 | 0.2029 | 0.0336 | 482723 | 4243 | 2517 | common | fungivorous | moist and cold | deciduous | 6.5000 |
| Bolitophagus reticulatus | Tenebrionidae | KF | KFe | Managed | 53.3391 | 23.3275 | 0.0020 | 0.2134 | 0.2042 | 0.0369 | 484203 | 4243 | 2525 | common | fungivorous | moist and cold | deciduous | 6.5000 |
| Bolitophagus reticulatus | Tenebrionidae | AF | AFa | Managed | 54.0433 | 23.4698 | 0.0024 | 0.2694 | 0.2375 | 0.0125 | 486478 | 4235 | 2724 | common | fungivorous | moist and cold | deciduous | 6.5000 |
| Boros schneideri | Boridae | BF | BFc | Primeval | 52.6684 | 23.7689 | 0.0024 | 0.1782 | 0.1881 | 0.0851 | 978627 | 10908 | 6088 | rare | invertivorous | moist and cold | coniferous | 12.5000 |
| Boros schneideri | Boridae | HM | HMe | Primeval | 51.0434 | 20.6831 | 0.0019 | 0.1627 | 0.1541 | 0.0232 | 977004 | 10902 | 4647 | rare | invertivorous | moist and cold | coniferous | 12.5000 |
| Boros schneideri | Boridae | CM | CMd | Primeval | 49.4301 | 22.6204 | 0.0023 | 0.1742 | 0.1826 | 0.0767 | 977205 | 10905 | 5675 | rare | invertivorous | moist and cold | coniferous | 12.5000 |
| Boros schneideri | Boridae | KF | KFe | Managed | 53.1461 | 23.6814 | 0.0021 | 0.1719 | 0.1638 | 0.0311 | 976628 | 10902 | 4979 | rare | invertivorous | moist and cold | coniferous | 12.5000 |
| Boros schneideri | Boridae | AF | AFa | Managed | 53.8614 | 23.3308 | 0.0022 | 0.1614 | 0.1721 | 0.0709 | 977260 | 10908 | 5471 | rare | invertivorous | moist and cold | coniferous | 12.5000 |
| Cucujus cinnaberinus | Cucujidae | BF | BFc | Primeval | 52.6684 | 23.7689 | 0.0019 | 0.2157 | 0.1677 | -0.0406 | 812685 | 8043 | 3528 | rare | invertivorous | moist and cold | various | 13 |
| Cucujus cinnaberinus | Cucujidae | HM | HMe | Primeval | 50.9653 | 20.4835 | 0.0022 | 0.2255 | 0.1996 | 0.0034 | 818291 | 8099 | 4604 | rare | invertivorous | moist and cold | various | 13 |
| Cucujus cinnaberinus | Cucujidae | CM | CMg | Primeval | 49.5765 | 22.4696 | 0.0025 | 0.2223 | 0.2265 | 0.0672 | 818228 | 8100 | 5384 | rare | invertivorous | moist and cold | various | 13 |
| Cucujus cinnaberinus | Cucujidae | KF | KFa | Managed | 53.2032 | 23.5788 | 0.0022 | 0.2052 | 0.1991 | 0.0447 | 818338 | 8098 | 4600 | rare | invertivorous | moist and cold | various | 13 |
| Cucujus cinnaberinus | Cucujidae | AF | AFc | Managed | 53.8700 | 23.3624 | 0.0023 | 0.2404 | 0.2060 | -0.0133 | 813968 | 8066 | 4666 | rare | invertivorous | moist and cold | various | 13 |
| Cucujus cinnaberinus | Cucujidae | OF | OFb | Managed | 50.0110 | 18.2659 | 0.0023 | 0.2030 | 0.2082 | 0.0651 | 816511 | 8097 | 4737 | rare | invertivorous | moist and cold | various | 13 |
| Cucujus haematodes | Cucujidae | AF | AFb | Managed | 53.8124 | 23.4517 | 0.0032 | 0.2227 | 0.2339 | 0.0964 | 962285 | 11541 | 7737 | rare | invertivorous | moist and cold | various | 15 |
| Cucujus haematodes | Cucujidae | BF | BFa | Primeval | 52.6684 | 23.7689 | 0.0030 | 0.2098 | 0.2263 | 0.0978 | 972396 | 11569 | 7734 | rare | invertivorous | moist and cold | various | 15 |
| Cucujus haematodes | Cucujidae | BF | BFb | Primeval | 52.6334 | 23.7286 | 0.0031 | 0.2024 | 0.2345 | 0.1344 | 990992 | 11569 | 8040 | rare | invertivorous | moist and cold | various | 15 |
| Cucujus haematodes | Cucujidae | HM | HMa | Primeval | 51.0512 | 20.7985 | 0.0031 | 0.1987 | 0.2308 | 0.1326 | 965291 | 11569 | 7948 | rare | invertivorous | moist and cold | various | 15 |
| Cucujus haematodes | Cucujidae | HM | HMb | Primeval | 51.0585 | 20.7033 | 0.0028 | 0.2033 | 0.2116 | 0.0769 | 971806 | 11567 | 7129 | rare | invertivorous | moist and cold | various | 15 |
| Chalcophora mariana | Buprestidae | SF | SFa | Managed | 51.3248 | 15.4453 | 0.0028 | 0.2053 | 0.2080 | 0.0661 | 5564717 | 66926 | 41965 | pest | xylovorous | dry and warm | coniferous | 31.5000 |
| Chalcophora mariana | Buprestidae | SF | SFb | Managed | 51.5012 | 15.6974 | 0.0029 | 0.1945 | 0.2156 | 0.1055 | 5567553 | 66929 | 44061 | pest | xylovorous | dry and warm | coniferous | 31.5000 |
| Chalcophora mariana | Buprestidae | DB | DBa | Managed | 51.5696 | 17.2649 | 0.0028 | 0.1963 | 0.2110 | 0.0894 | 5586249 | 66928 | 43107 | pest | xylovorous | dry and warm | coniferous | 31.5000 |
| Chalcophora mariana | Buprestidae | DB | DBb | Managed | 51.5272 | 17.2574 | 0.0030 | 0.2157 | 0.2176 | 0.0757 | 5558736 | 66884 | 41750 | pest | xylovorous | dry and warm | coniferous | 31.5000 |
| Chalcophora mariana | Buprestidae | KF | KFa | Managed | 53.2538 | 23.7603 | 0.0029 | 0.1990 | 0.2068 | 0.0846 | 5557123 | 66896 | 38694 | pest | xylovorous | dry and warm | coniferous | 31.5000 |
| Chalcophora mariana | Buprestidae | AF | AFa | Managed | 53.8741 | 23.3067 | 0.0028 | 0.1953 | 0.2091 | 0.0904 | 5560437 | 66926 | 41143 | pest | xylovorous | dry and warm | coniferous | 31.5000 |
| Diaperis boleti | Tenebrionidae | CM | CMa | Primeval | 49.7734 | 22.6787 | 0.0032 | 0.2058 | 0.2275 | 0.1129 | 1669023 | 20987 | 14165 | common | fungivorous | moist and cold | deciduous | 7 |
| Diaperis boleti | Tenebrionidae | CM | CMb | Primeval | 49.5509 | 22.6025 | 0.0031 | 0.1907 | 0.2208 | 0.1262 | 1668402 | 20984 | 13730 | common | fungivorous | moist and cold | deciduous | 7 |
| Diaperis boleti | Tenebrionidae | BF | BFa | Primeval | 52.7230 | 23.8536 | 0.0031 | 0.1882 | 0.2195 | 0.1290 | 1668735 | 20987 | 13620 | common | fungivorous | moist and cold | deciduous | 7 |
| Diaperis boleti | Tenebrionidae | HM | HMa | Primeval | 51.0591 | 20.7015 | 0.0033 | 0.2001 | 0.2344 | 0.1381 | 1667035 | 20984 | 14525 | common | fungivorous | moist and cold | deciduous | 7 |
| Diaperis boleti | Tenebrionidae | HM | HMb | Primeval | 51.0585 | 20.7033 | 0.0033 | 0.2036 | 0.2370 | 0.1390 | 1666886 | 20986 | 14642 | common | fungivorous | moist and cold | deciduous | 7 |
| Elateroides dermestoides | Lymexylidae | AF | AFa | Managed | 54.0029 | 23.4686 | 0.0032 | 0.1755 | 0.2261 | 0.1615 | 992483 | 12412 | 8228 | pest | xylovorous | moist and cold | various | 9 |
| Elateroides dermestoides | Lymexylidae | CM | CMa | Primeval | 49.2534 | 22.5611 | 0.0027 | 0.1781 | 0.1927 | 0.0852 | 992580 | 12412 | 7317 | pest | xylovorous | moist and cold | various | 9 |
| Elateroides dermestoides | Lymexylidae | CM | CMb | Primeval | 49.4242 | 22.6584 | 0.0028 | 0.1839 | 0.1976 | 0.0844 | 992489 | 12412 | 7528 | pest | xylovorous | moist and cold | various | 9 |
| Elater ferrugineus | Elateridae | BF | BFb | Primeval | 52.8176 | 23.7593 | 0.0030 | 0.3474 | 0.2814 | -0.0500 | 1644540 | 15761 | 11257 | rare | invertivorous | moist and cold | deciduous | 20.5000 |
| Elater ferrugineus | Elateridae | HM | HMa | Primeval | 51.0591 | 20.7015 | 0.0032 | 0.3187 | 0.2937 | 0.0329 | 1644301 | 15766 | 11944 | rare | invertivorous | moist and cold | deciduous | 20.5000 |
| Elater ferrugineus | Elateridae | DB | DBl | Managed | 51.4786 | 16.9243 | 0.0033 | 0.3147 | 0.3032 | 0.0686 | 1641138 | 15763 | 12295 | rare | invertivorous | moist and cold | deciduous | 20.5000 |
| Elater ferrugineus | Elateridae | OF | OFc | Managed | 50.9421 | 17.3905 | 0.0030 | 0.2606 | 0.2822 | 0.1192 | 1642493 | 15765 | 11743 | rare | invertivorous | moist and cold | deciduous | 20.5000 |
| Elater ferrugineus | Elateridae | BF | BFh | Primeval | 52.6932 | 23.8137 | 0.0031 | 0.3056 | 0.2887 | 0.0510 | 1645687 | 15766 | 12365 | rare | invertivorous | moist and cold | deciduous | 20.5000 |
| Elater ferrugineus | Elateridae | OF | OFa | Managed | 50.9463 | 17.3281 | 0.0029 | 0.2346 | 0.2723 | 0.1534 | 1641108 | 15765 | 11263 | rare | invertivorous | moist and cold | deciduous | 20.5000 |
| Ips acuminatus | Curculionidae | HM | HMb | Primeval | 50.8856 | 21.0789 | 0.0027 | 0.1265 | 0.1661 | 0.1375 | 563184 | 8066 | 4104 | pest | xylovorous | dry and warm | coniferous | 2.8500 |
| Ips acuminatus | Curculionidae | HM | HMc | Primeval | 50.8858 | 21.0717 | 0.0026 | 0.1183 | 0.1618 | 0.1491 | 564178 | 8067 | 3948 | pest | xylovorous | dry and warm | coniferous | 2.8500 |
| Ips acuminatus | Curculionidae | HM | HMd | Primeval | 50.8677 | 21.0635 | 0.0026 | 0.1190 | 0.1580 | 0.1405 | 563327 | 8058 | 3591 | pest | xylovorous | dry and warm | coniferous | 2.8500 |
| Ips acuminatus | Curculionidae | KF | KFa | Managed | 53.1451 | 23.7970 | 0.0024 | 0.0998 | 0.1499 | 0.1614 | 562859 | 8055 | 3565 | pest | xylovorous | dry and warm | coniferous | 2.8500 |
| Ips acuminatus | Curculionidae | KF | KFf | Managed | 53.1514 | 23.8218 | 0.0029 | 0.1058 | 0.1755 | 0.2137 | 561709 | 8043 | 4212 | pest | xylovorous | dry and warm | coniferous | 2.8500 |
| Ips acuminatus | Curculionidae | AF | AFb | Managed | 53.9255 | 23.2311 | 0.0019 | 0.1106 | 0.1181 | 0.0656 | 560817 | 8039 | 2748 | pest | xylovorous | dry and warm | coniferous | 2.8500 |
| Ips sexdentatus | Curculionidae | BF | BFd | Primeval | 52.6376 | 23.5500 | 0.0040 | 0.1053 | 0.1564 | 0.1629 | 554961 | 12536 | 6095 | pest | xylovorous | dry and warm | coniferous | 6.5000 |
| Ips sexdentatus | Curculionidae | BF | BFe | Primeval | 52.6455 | 23.7008 | 0.0037 | 0.1007 | 0.1409 | 0.1313 | 554672 | 12531 | 5162 | pest | xylovorous | dry and warm | coniferous | 6.5000 |
| Ips sexdentatus | Curculionidae | HM | HMe | Primeval | 50.8677 | 21.0635 | 0.0040 | 0.1107 | 0.1567 | 0.1540 | 553606 | 12536 | 6212 | pest | xylovorous | dry and warm | coniferous | 6.5000 |
| Ips sexdentatus | Curculionidae | KF | KFa | Managed | 53.1451 | 23.7970 | 0.0042 | 0.1179 | 0.1634 | 0.1524 | 553117 | 12536 | 6434 | pest | xylovorous | dry and warm | coniferous | 6.5000 |
| Ips sexdentatus | Curculionidae | KF | KFf | Managed | 53.1514 | 23.8218 | 0.0038 | 0.1150 | 0.1504 | 0.1267 | 554641 | 12529 | 5940 | pest | xylovorous | dry and warm | coniferous | 6.5000 |
| Ips sexdentatus | Curculionidae | AF | AFc | Managed | 53.9255 | 23.2311 | 0.0038 | 0.0974 | 0.1479 | 0.1621 | 553438 | 12528 | 5457 | pest | xylovorous | dry and warm | coniferous | 6.5000 |
| Ips typographus | Curculionidae | BF | BFh | Primeval | 52.6376 | 23.5500 | 0.0033 | 0.1614 | 0.1870 | 0.1119 | 342428 | 5360 | 3058 | pest | xylovorous | moist and cold | coniferous | 4.8500 |
| Ips typographus | Curculionidae | HM | HMb | Primeval | 51.0465 | 20.6699 | 0.0031 | 0.1331 | 0.1737 | 0.1451 | 342193 | 5353 | 2839 | pest | xylovorous | moist and cold | coniferous | 4.8500 |
| Ips typographus | Curculionidae | CM | CMd | Primeval | 49.4305 | 22.6481 | 0.0031 | 0.1422 | 0.1746 | 0.1244 | 342546 | 5355 | 2883 | pest | xylovorous | moist and cold | coniferous | 4.8500 |
| Ips typographus | Curculionidae | KF | KFc | Managed | 53.3456 | 23.3134 | 0.0034 | 0.1592 | 0.1916 | 0.1263 | 342706 | 5361 | 3130 | pest | xylovorous | moist and cold | coniferous | 4.8500 |
| Ips typographus | Curculionidae | AF | AFd | Managed | 53.8491 | 23.3644 | 0.0032 | 0.1693 | 0.1833 | 0.0858 | 341706 | 5357 | 2974 | pest | xylovorous | moist and cold | coniferous | 4.8500 |
| Ips typographus | Curculionidae | SF | SFa | Managed | 51.3391 | 15.2616 | 0.0031 | 0.1660 | 0.1706 | 0.0704 | 341451 | 5346 | 2730 | pest | xylovorous | moist and cold | coniferous | 4.8500 |
| Monochamus galloprovincialis | Cerambycidae | SF | SFa | Managed | 51.5249 | 15.7700 | 0.0038 | 0.1368 | 0.1743 | 0.1356 | 2608622 | 50483 | 28006 | pest | xylovorous | dry and warm | coniferous | 18.5000 |
| Monochamus galloprovincialis | Cerambycidae | SF | SFb | Managed | 51.5058 | 15.6849 | 0.0036 | 0.1344 | 0.1634 | 0.1131 | 2608920 | 50483 | 26268 | pest | xylovorous | dry and warm | coniferous | 18.5000 |
| Monochamus galloprovincialis | Cerambycidae | CM | CMa | Primeval | 49.5479 | 22.5257 | 0.0037 | 0.1434 | 0.1675 | 0.1035 | 2608306 | 50480 | 26485 | pest | xylovorous | dry and warm | coniferous | 18.5000 |
| Monochamus galloprovincialis | Cerambycidae | HM | HMa | Primeval | 51.0807 | 20.7713 | 0.0037 | 0.1304 | 0.1693 | 0.1361 | 2611964 | 50484 | 26088 | pest | xylovorous | dry and warm | coniferous | 18.5000 |
| Monochamus galloprovincialis | Cerambycidae | AF | AFa | Managed | 53.8741 | 23.3067 | 0.0037 | 0.1401 | 0.1659 | 0.1147 | 2605446 | 50426 | 25015 | pest | xylovorous | dry and warm | coniferous | 18.5000 |
| Monochamus galloprovincialis | Cerambycidae | BF | BFa | Primeval | 52.6229 | 23.5405 | 0.0031 | 0.1384 | 0.1418 | 0.0527 | 2602150 | 50455 | 20196 | pest | xylovorous | dry and warm | coniferous | 18.5000 |
| Monochamus galloprovincialis | Cerambycidae | KF | KFa | Managed | 53.1705 | 23.8189 | 0.0037 | 0.1377 | 0.1722 | 0.1273 | 2609741 | 50484 | 27186 | pest | xylovorous | dry and warm | coniferous | 18.5000 |
| Neomida haemorrhoidalis | Tenebrionidae | SF | SFb | Managed | 51.5372 | 17.3952 | 0.0021 | 0.2113 | 0.1924 | 0.0208 | 1591986 | 15208 | 8063 | rare | fungivorous | moist and cold | deciduous | 5.7500 |
| Neomida haemorrhoidalis | Tenebrionidae | DB | DBd | Managed | 51.5055 | 15.6847 | 0.0022 | 0.2060 | 0.2023 | 0.0554 | 1593409 | 15222 | 8844 | rare | fungivorous | moist and cold | deciduous | 5.7500 |
| Neomida haemorrhoidalis | Tenebrionidae | BF | BFf | Primeval | 52.7863 | 23.7127 | 0.0021 | 0.1817 | 0.1962 | 0.0858 | 1592871 | 15235 | 9156 | rare | fungivorous | moist and cold | deciduous | 5.7500 |
| Neomida haemorrhoidalis | Tenebrionidae | HM | HMb | Primeval | 50.8853 | 21.1053 | 0.0020 | 0.1797 | 0.1883 | 0.0702 | 1594210 | 15234 | 8531 | rare | fungivorous | moist and cold | deciduous | 5.7500 |
| Neomida haemorrhoidalis | Tenebrionidae | CM | CMc | Primeval | 49.5325 | 22.5684 | 0.0021 | 0.1825 | 0.1996 | 0.0912 | 1598147 | 15235 | 9349 | rare | fungivorous | moist and cold | deciduous | 5.7500 |
| Neomida haemorrhoidalis | Tenebrionidae | KF | KFa | Managed | 53.2077 | 23.5815 | 0.0022 | 0.2075 | 0.1996 | 0.0390 | 1594280 | 15235 | 9216 | rare | fungivorous | moist and cold | deciduous | 5.7500 |
| Neomida haemorrhoidalis | Tenebrionidae | AF | AFd | Managed | 53.9265 | 23.4242 | 0.0021 | 0.1988 | 0.1921 | 0.0435 | 1590429 | 15220 | 8760 | rare | fungivorous | moist and cold | deciduous | 5.7500 |
| Osmoderma barnabita | Scarabaeidae | SF | SFa | Managed | 51.5012 | 15.6974 | 0.0024 | 0.2596 | 0.2442 | 0.0603 | 921185 | 7798 | 5288 | rare | xylovorous | moist and cold | deciduous | 27 |
| Osmoderma barnabita | Scarabaeidae | DB | DBa | Managed | 51.4786 | 16.9243 | 0.0029 | 0.3895 | 0.3050 | -0.0574 | 935978 | 7896 | 5778 | rare | xylovorous | moist and cold | deciduous | 27 |
| Osmoderma barnabita | Scarabaeidae | DB | DBb | Managed | 51.5272 | 17.2574 | 0.0024 | 0.3152 | 0.2547 | -0.0410 | 936547 | 7895 | 5541 | rare | xylovorous | moist and cold | deciduous | 27 |
| Osmoderma barnabita | Scarabaeidae | OF | OFa | Managed | 50.9421 | 17.3905 | 0.0022 | 0.2590 | 0.2256 | -0.0007 | 935807 | 7884 | 4642 | rare | xylovorous | moist and cold | deciduous | 27 |
| Osmoderma barnabita | Scarabaeidae | AF | AFa | Managed | 54.0029 | 23.4686 | 0.0004 | 0.0411 | 0.0408 | 0.0112 | 932545 | 7890 | 927 | rare | xylovorous | moist and cold | deciduous | 27 |
| Osmoderma barnabita | Scarabaeidae | BF | BFa | Primeval | 52.8176 | 23.7593 | 0.0005 | 0.0648 | 0.0540 | -0.0085 | 935314 | 7898 | 1414 | rare | xylovorous | moist and cold | deciduous | 27 |
| Osmoderma barnabita | Scarabaeidae | BF | BFb | Primeval | 52.6932 | 23.8137 | 0.0005 | 0.0631 | 0.0486 | -0.0173 | 934941 | 7898 | 1079 | rare | xylovorous | moist and cold | deciduous | 27 |
| Osmoderma barnabita | Scarabaeidae | CM | CMa | Primeval | 49.7734 | 22.6787 | 0.0019 | 0.3059 | 0.1845 | -0.1304 | 934411 | 7880 | 3175 | rare | xylovorous | moist and cold | deciduous | 27 |
| Osmoderma barnabita | Scarabaeidae | HM | HMa | Primeval | 51.0591 | 20.7015 | 0.0019 | 0.2920 | 0.2059 | -0.1085 | 936866 | 7898 | 3649 | rare | xylovorous | moist and cold | deciduous | 27 |
| Osmoderma barnabita | Scarabaeidae | HM | HMb | Primeval | 50.8825 | 21.1059 | 0.0013 | 0.1991 | 0.1373 | -0.0773 | 935485 | 7896 | 2535 | rare | xylovorous | moist and cold | deciduous | 27 |
| Phaenops cyanea | Buprestidae | SF | SFd | Managed | 51.4178 | 15.8514 | 0.0027 | 0.2433 | 0.2198 | 0.0193 | 3479542 | 38333 | 23820 | pest | xylovorous | dry and warm | coniferous | 9.5000 |
| Phaenops cyanea | Buprestidae | BF | BFc | Primeval | 52.6455 | 23.7008 | 0.0025 | 0.2073 | 0.2025 | 0.0476 | 3475768 | 38341 | 23670 | pest | xylovorous | dry and warm | coniferous | 9.5000 |
| Phaenops cyanea | Buprestidae | HM | HMa | Primeval | 50.8553 | 20.5904 | 0.0028 | 0.2340 | 0.2268 | 0.0584 | 3477909 | 38338 | 24708 | pest | xylovorous | dry and warm | coniferous | 9.5000 |
| Phaenops cyanea | Buprestidae | CM | CMa | Primeval | 49.6459 | 22.5059 | 0.0026 | 0.2340 | 0.2073 | 0.0062 | 3478924 | 38338 | 23333 | pest | xylovorous | dry and warm | coniferous | 9.5000 |
| Phaenops cyanea | Buprestidae | KF | KFb | Managed | 53.1396 | 23.7401 | 0.0026 | 0.2069 | 0.2133 | 0.0734 | 3471842 | 38337 | 24367 | pest | xylovorous | dry and warm | coniferous | 9.5000 |
| Phaenops cyanea | Buprestidae | AF | AFc | Managed | 53.8287 | 23.4851 | 0.0025 | 0.2117 | 0.1968 | 0.0418 | 3287112 | 35963 | 20519 | pest | xylovorous | dry and warm | coniferous | 9.5000 |
| Pytho depressus | Pythidae | AF | AFb | Managed | 53.8248 | 23.3389 | 0.0019 | 0.1344 | 0.1058 | -0.0213 | 269393 | 4200 | 1206 | common | invertivorous | moist and cold | various | 11.7500 |
| Pytho depressus | Pythidae | KF | KFa | Managed | 53.1269 | 23.7816 | 0.0022 | 0.1068 | 0.1238 | 0.0723 | 275232 | 4219 | 1532 | common | invertivorous | moist and cold | various | 11.7500 |
| Pytho depressus | Pythidae | KF | KFb | Managed | 53.2042 | 23.5831 | 0.0023 | 0.1304 | 0.1285 | 0.0363 | 269470 | 4221 | 1621 | common | invertivorous | moist and cold | various | 11.7500 |
| Pytho depressus | Pythidae | CM | CMa | Primeval | 49.5912 | 22.6382 | 0.0023 | 0.1200 | 0.1272 | 0.0570 | 266401 | 4217 | 1595 | common | invertivorous | moist and cold | various | 11.7500 |
| Pytho depressus | Pythidae | HM | HMa | Primeval | 50.9657 | 20.4750 | 0.0026 | 0.1337 | 0.1432 | 0.0630 | 265544 | 4221 | 1885 | common | invertivorous | moist and cold | various | 11.7500 |
| Pytho depressus | Pythidae | BF | BFa | Primeval | 52.7076 | 23.7745 | 0.0026 | 0.1293 | 0.1487 | 0.0886 | 266251 | 4221 | 2071 | common | invertivorous | moist and cold | various | 11.7500 |
| Pytho depressus | Pythidae | AF | AFa | Managed | 53.8248 | 23.3389 | 0.0026 | 0.1326 | 0.1482 | 0.0830 | 267629 | 4221 | 2021 | common | invertivorous | moist and cold | various | 11.7500 |
| Pyrochroa coccinea | Pyrochroidae | SF | SFa | Managed | 51.5012 | 15.6974 | 0.0020 | 0.2318 | 0.2219 | 0.0397 | 734679 | 5916 | 3800 | common | invertivorous | moist and cold | deciduous | 16 |
| Pyrochroa coccinea | Pyrochroidae | OF | OFa | Managed | 51.5032 | 15.6916 | 0.0024 | 0.3125 | 0.2566 | -0.0372 | 733803 | 5909 | 4132 | common | invertivorous | moist and cold | deciduous | 16 |
| Pyrochroa coccinea | Pyrochroidae | CM | CMa | Primeval | 49.5910 | 22.6395 | 0.0023 | 0.2923 | 0.2418 | -0.0130 | 735255 | 5907 | 3719 | common | invertivorous | moist and cold | deciduous | 16 |
| Pyrochroa coccinea | Pyrochroidae | HM | HMa | Primeval | 51.0764 | 20.7171 | 0.0021 | 0.2767 | 0.2311 | -0.0288 | 734120 | 5915 | 3791 | common | invertivorous | moist and cold | deciduous | 16 |
| Pyrochroa coccinea | Pyrochroidae | BF | BFa | Primeval | 52.7217 | 23.9162 | 0.0020 | 0.2489 | 0.2160 | -0.0073 | 734542 | 5916 | 3656 | common | invertivorous | moist and cold | deciduous | 16 |
| Pyrochroa coccinea | Pyrochroidae | KF | KFa | Managed | 53.3425 | 23.3002 | 0.0020 | 0.2363 | 0.2203 | 0.0303 | 735157 | 5916 | 3717 | common | invertivorous | moist and cold | deciduous | 16 |
| Pyrochroa coccinea | Pyrochroidae | AF | AFa | Managed | 54.0612 | 23.3617 | 0.0019 | 0.2249 | 0.2156 | 0.0401 | 734525 | 5916 | 3643 | common | invertivorous | moist and cold | deciduous | 16 |
| Rhagium inquisitor | Cerambycidae | BF | BFc | Primeval | 52.6292 | 23.7131 | 0.0038 | 0.1233 | 0.1514 | 0.1041 | 732634 | 16340 | 7527 | common | xylovorous | moist and cold | coniferous | 15.5000 |
| Rhagium inquisitor | Cerambycidae | HM | HMd | Primeval | 50.9666 | 20.4791 | 0.0042 | 0.1188 | 0.1670 | 0.1571 | 732230 | 16337 | 8494 | common | xylovorous | moist and cold | coniferous | 15.5000 |
| Rhagium inquisitor | Cerambycidae | CM | CMb | Primeval | 49.6839 | 22.6034 | 0.0043 | 0.1326 | 0.1674 | 0.1304 | 730239 | 16343 | 8257 | common | xylovorous | moist and cold | coniferous | 15.5000 |
| Rhagium inquisitor | Cerambycidae | KF | KFb | Managed | 53.4106 | 23.3800 | 0.0043 | 0.1320 | 0.1682 | 0.1319 | 729771 | 16337 | 8272 | common | xylovorous | moist and cold | coniferous | 15.5000 |
| Rhagium inquisitor | Cerambycidae | AF | AFa | Managed | 53.8725 | 23.3056 | 0.0044 | 0.1310 | 0.1723 | 0.1426 | 728388 | 16346 | 8564 | common | xylovorous | moist and cold | coniferous | 15.5000 |
| Rhagium inquisitor | Cerambycidae | DB | DBb | Managed | 51.4385 | 17.2205 | 0.0046 | 0.1332 | 0.1835 | 0.1662 | 729938 | 16344 | 9396 | common | xylovorous | moist and cold | coniferous | 15.5000 |
| Rhysodes sulcatus | Carabidae | KF | KFa | Managed | 53.2042 | 23.5831 | 0.0021 | 0.5206 | 0.3046 | -0.3205 | 88562 | 540 | 378 | rare | xylovorous | moist and cold | various | 7 |
| Rhysodes sulcatus | Carabidae | CM | CMa | Primeval | 49.5479 | 22.5257 | 0.0023 | 0.5701 | 0.3317 | -0.3586 | 88597 | 540 | 410 | rare | xylovorous | moist and cold | various | 7 |
| Rhysodes sulcatus | Carabidae | CM | CMb | Primeval | 49.2534 | 22.5611 | 0.0020 | 0.4967 | 0.2938 | -0.2905 | 88583 | 540 | 371 | rare | xylovorous | moist and cold | various | 7 |
| Rhysodes sulcatus | Carabidae | HM | HMa | Primeval | 51.0534 | 20.6832 | 0.0020 | 0.4740 | 0.2875 | -0.2509 | 88644 | 540 | 361 | rare | xylovorous | moist and cold | various | 7 |
| Spondylis buprestoides | Cerambycidae | SF | SFa | Managed | 51.3274 | 15.4430 | 0.0029 | 0.1933 | 0.2008 | 0.0701 | 4087571 | 52548 | 32959 | common | xylovorous | dry and warm | coniferous | 18 |
| Spondylis buprestoides | Cerambycidae | SF | SFb | Managed | 51.5012 | 15.6974 | 0.0029 | 0.1946 | 0.2016 | 0.0688 | 4090006 | 52548 | 33267 | common | xylovorous | dry and warm | coniferous | 18 |
| Spondylis buprestoides | Cerambycidae | AF | AFa | Managed | 53.8741 | 23.3067 | 0.0029 | 0.1927 | 0.2010 | 0.0729 | 4093279 | 52547 | 33766 | common | xylovorous | dry and warm | coniferous | 18 |
| Spondylis buprestoides | Cerambycidae | KF | KFa | Managed | 53.1976 | 23.7312 | 0.0030 | 0.2118 | 0.2044 | 0.0530 | 4087935 | 52499 | 31439 | common | xylovorous | dry and warm | coniferous | 18 |
| Spondylis buprestoides | Cerambycidae | BF | BFa | Primeval | 52.6229 | 23.5405 | 0.0029 | 0.1997 | 0.2024 | 0.0646 | 4089707 | 52545 | 32553 | common | xylovorous | dry and warm | coniferous | 18 |
| Spondylis buprestoides | Cerambycidae | HM | HMa | Primeval | 51.0534 | 20.6832 | 0.0028 | 0.1936 | 0.1881 | 0.0523 | 4085567 | 52513 | 28189 | common | xylovorous | dry and warm | coniferous | 18 |
| Spondylis buprestoides | Cerambycidae | CM | CMa | Primeval | 49.6744 | 22.4588 | 0.0029 | 0.2019 | 0.2053 | 0.0622 | 4088396 | 52549 | 33575 | common | xylovorous | dry and warm | coniferous | 18 |
| Saperda scalaris | Cerambycidae | BF | BFa | Primeval | 52.7160 | 23.6067 | 0.0035 | 0.1865 | 0.2152 | 0.1206 | 1191830 | 17219 | 11262 | pest | xylovorous | moist and cold | deciduous | 15 |
| Saperda scalaris | Cerambycidae | KF | KFa | Managed | 53.1269 | 23.7816 | 0.0037 | 0.2022 | 0.2258 | 0.1099 | 1192688 | 17219 | 11720 | pest | xylovorous | moist and cold | deciduous | 15 |
| Saperda scalaris | Cerambycidae | AF | AFa | Managed | 53.8124 | 23.4517 | 0.0038 | 0.2379 | 0.2326 | 0.0571 | 1190513 | 17219 | 11929 | pest | xylovorous | moist and cold | deciduous | 15 |
| Scolytus intricatus | Curculionidae | HM | HMa | Primeval | 51.0534 | 20.6832 | 0.0028 | 0.2119 | 0.2382 | 0.1233 | 906870 | 9553 | 6743 | pest | xylovorous | dry and warm | deciduous | 3 |
| Scolytus intricatus | Curculionidae | BF | BFa | Primeval | 52.6229 | 23.5405 | 0.0026 | 0.2245 | 0.2095 | 0.0431 | 905671 | 9548 | 5526 | pest | xylovorous | dry and warm | deciduous | 3 |
| Scolytus intricatus | Curculionidae | KF | KFa | Managed | 53.4115 | 23.3748 | 0.0027 | 0.2279 | 0.2238 | 0.0555 | 906769 | 9554 | 6211 | pest | xylovorous | dry and warm | deciduous | 3 |
| Scolytus intricatus | Curculionidae | AF | AFa | Managed | 54.0029 | 23.4686 | 0.0027 | 0.2016 | 0.2313 | 0.1266 | 907005 | 9554 | 6464 | pest | xylovorous | dry and warm | deciduous | 3 |
| Stenurella melanura | Cerambycidae | BF | BFd | Primeval | 52.6596 | 23.6211 | 0.0057 | 0.0880 | 0.1306 | 0.1362 | 1487513 | 57466 | 23420 | common | xylovorous | moist and cold | various | 7.5000 |
| Stenurella melanura | Cerambycidae | HM | HMe | Primeval | 51.0607 | 20.6968 | 0.0064 | 0.0935 | 0.1461 | 0.1646 | 1486037 | 57467 | 26389 | common | xylovorous | moist and cold | various | 7.5000 |
| Stenurella melanura | Cerambycidae | CM | CMa | Primeval | 49.7993 | 22.6629 | 0.0061 | 0.0892 | 0.1406 | 0.1599 | 1484420 | 57449 | 25234 | common | xylovorous | moist and cold | various | 7.5000 |
| Stenurella melanura | Cerambycidae | KF | KFb | Managed | 53.3001 | 23.1263 | 0.0062 | 0.0896 | 0.1429 | 0.1658 | 1484423 | 57456 | 25918 | common | xylovorous | moist and cold | various | 7.5000 |
| Stenurella melanura | Cerambycidae | AF | AFc | Managed | 53.8741 | 23.3486 | 0.0061 | 0.0901 | 0.1409 | 0.1590 | 1484858 | 57455 | 25601 | common | xylovorous | moist and cold | various | 7.5000 |
| Stenurella melanura | Cerambycidae | OF | OFb | Managed | 50.9434 | 17.3930 | 0.0059 | 0.0848 | 0.1340 | 0.1522 | 1478333 | 57327 | 23058 | common | xylovorous | moist and cold | various | 7.5000 |
| Schizotus pectinicornis | Pyrochroidae | OF | OFa | Managed | 51.5032 | 15.6916 | 0.0019 | 0.1561 | 0.1491 | 0.0342 | 463955 | 5061 | 2061 | common | invertivorous | moist and cold | deciduous | 8.5000 |
| Schizotus pectinicornis | Pyrochroidae | HM | HMb | Primeval | 51.0764 | 20.7171 | 0.0022 | 0.1760 | 0.1800 | 0.0544 | 482233 | 5234 | 2728 | common | invertivorous | moist and cold | deciduous | 8.5000 |
| Schizotus pectinicornis | Pyrochroidae | CM | CMd | Primeval | 49.5910 | 22.6395 | 0.0024 | 0.1865 | 0.1956 | 0.0726 | 482468 | 5234 | 2976 | common | invertivorous | moist and cold | deciduous | 8.5000 |
| Schizotus pectinicornis | Pyrochroidae | BF | BFe | Primeval | 52.7217 | 23.9162 | 0.0021 | 0.1747 | 0.1749 | 0.0476 | 482437 | 5234 | 2601 | common | invertivorous | moist and cold | deciduous | 8.5000 |
| Schizotus pectinicornis | Pyrochroidae | KF | KFc | Managed | 53.3425 | 23.3002 | 0.0023 | 0.1980 | 0.1838 | 0.0279 | 481726 | 5229 | 2626 | common | invertivorous | moist and cold | deciduous | 8.5000 |
| Schizotus pectinicornis | Pyrochroidae | AF | AFd | Managed | 54.0612 | 23.3617 | 0.0021 | 0.1740 | 0.1716 | 0.0405 | 481499 | 5232 | 2574 | common | invertivorous | moist and cold | deciduous | 8.5000 |
| Thanasimus formicarius | Cleridae | HM | HMd | Primeval | 51.0534 | 20.7037 | 0.0040 | 0.1430 | 0.1759 | 0.1267 | 5188533 | 104610 | 57689 | common | invertivorous | dry and warm | coniferous | 8.5000 |
| Thanasimus formicarius | Cleridae | KF | KFb | Managed | 53.1540 | 23.8888 | 0.0044 | 0.1537 | 0.1926 | 0.1417 | 5179910 | 104622 | 62221 | common | invertivorous | dry and warm | coniferous | 8.5000 |
| Thanasimus formicarius | Cleridae | AF | AFh | Managed | 53.9609 | 23.4805 | 0.0043 | 0.1447 | 0.1898 | 0.1570 | 5188081 | 104565 | 61088 | common | invertivorous | dry and warm | coniferous | 8.5000 |
| Thanasimus formicarius | Cleridae | SF | SFb | Managed | 51.5205 | 15.7696 | 0.0043 | 0.1567 | 0.1897 | 0.1304 | 5177926 | 104618 | 61949 | common | invertivorous | dry and warm | coniferous | 8.5000 |
| Thanasimus formicarius | Cleridae | CM | CMd | Primeval | 49.6736 | 22.4557 | 0.0044 | 0.1625 | 0.1942 | 0.1283 | 5166529 | 104609 | 61733 | common | invertivorous | dry and warm | coniferous | 8.5000 |
| Uleiota planata | Silvanidae | AF | AFa | Managed | 53.8248 | 23.3389 | 0.0042 | 0.1468 | 0.1866 | 0.1421 | 2370375 | 47583 | 27599 | common | invertivorous | moist and cold | various | 5 |
| Uleiota planata | Silvanidae | CM | CMa | Primeval | 49.7734 | 22.6787 | 0.0041 | 0.1426 | 0.1796 | 0.1369 | 2371260 | 47583 | 27546 | common | invertivorous | moist and cold | various | 5 |
| Uleiota planata | Silvanidae | KF | KFa | Managed | 53.2538 | 23.7603 | 0.0041 | 0.1585 | 0.1806 | 0.1053 | 2362059 | 47456 | 26661 | common | invertivorous | moist and cold | various | 5 |
| Uleiota planata | Silvanidae | SF | SFa | Managed | 51.5272 | 17.2574 | 0.0045 | 0.1515 | 0.1985 | 0.1621 | 2367678 | 47570 | 29579 | common | invertivorous | moist and cold | various | 5 |

**Table S2. Species-level mean genomic-diversity metrics, sampling coverage and biological traits for the 26 saproxylic beetle species.**

| **Species** | **Ho** | **He** | **Pi** | **Fis** | **Family** | **Occurrence** | **Distribution** | **Conservation** | **Economic** | **Diet** | **Trophic guild** | **Microhabitat** | **Shade tolerance** | **Settlement** | **Tree type** | **Body size** | **BL min** | **BL max** | **BL mean** | **Dispersal ability** | **Voltinism** | **Total sites** | **Primeval** | **Reserves** | **Managed** | **Dispersal 2** | **Microhabitat 3** | **BL mean z** |
| --- | --- | --- | --- | --- | --- | --- | --- | --- | --- | --- | --- | --- | --- | --- | --- | --- | --- | --- | --- | --- | --- | --- | --- | --- | --- | --- | --- | --- |
| Bolitophagus reticulatus | 0.2267 | 0.2040 | 0.0020 | 0.0118 | Tenebrionidae | common | temperate | unprotected | neutral | fungivorous | mycetophagous | fungi | moist and cold | old | deciduous | small | 6 | 7 | 6.5000 | moderate | uni | 41 | 14 | 14 | 13 | low/moderate | fungi | -0.7359 |
| Boros schneideri | 0.1697 | 0.1721 | 0.0022 | 0.0574 | Boridae | rare | boreal | protected | neutral | invertivorous | predator & saprophagous | under bark | moist and cold | old | coniferous | medium | 11 | 14 | 12.5000 | moderate | uni | 32 | 18 | 4 | 10 | low/moderate | under bark | 0.1039 |
| Chalcophora mariana | 0.2010 | 0.2113 | 0.0029 | 0.0853 | Buprestidae | pest | temperate | unprotected | pest | xylovorous | xylophagous | wood | dry and warm | old | coniferous | large | 24 | 39 | 31.5000 | high | uni | 6 | 0 | 0 | 6 | high | wood/cavities | 2.7632 |
| Cucujus cinnaberinus | 0.2187 | 0.2012 | 0.0022 | 0.0211 | Cucujidae | rare | temperate | protected | neutral | invertivorous | predator & saprophagous | under bark | moist and cold | old | various | medium | 11 | 15 | 13 | moderate | uni | 40 | 17 | 8 | 15 | low/moderate | under bark | 0.1739 |
| Cucujus haematodes | 0.2074 | 0.2274 | 0.0031 | 0.1076 | Cucujidae | rare | boreal | protected | neutral | invertivorous | predator & saprophagous | under bark | moist and cold | old | various | medium | 13 | 17 | 15 | moderate | uni | 6 | 4 | 0 | 2 | low/moderate | under bark | 0.4538 |
| Diaperis boleti | 0.1977 | 0.2278 | 0.0032 | 0.1290 | Tenebrionidae | common | temperate | unprotected | neutral | fungivorous | mycetophagous | fungi | moist and cold | old | deciduous | small | 6 | 8 | 7 | moderate | uni | 5 | 5 | 0 | 0 | low/moderate | fungi | -0.6659 |
| Elater ferrugineus | 0.2969 | 0.2869 | 0.0031 | 0.0625 | Elateridae | rare | temperate | protected | neutral | invertivorous | predator | tree cavities | moist and cold | old | deciduous | large | 17 | 24 | 20.5000 | high | uni | 16 | 10 | 6 | 0 | high | wood/cavities | 1.2236 |
| Elateroides dermestoides | 0.1792 | 0.2055 | 0.0029 | 0.1104 | Lymexylidae | pest | temperate | unprotected | pest | xylovorous | xylophagous | wood | moist and cold | fresh | various | medium | 6 | 18 | 9 | moderate | uni | 4 | 3 | 0 | 1 | low/moderate | wood/cavities | -0.3860 |
| Ips acuminatus | 0.1134 | 0.1549 | 0.0025 | 0.1446 | Curculionidae | pest | temperate | unprotected | pest | xylovorous | cambiophagous | under bark | dry and warm | fresh | coniferous | small | 2.2000 | 3.5000 | 2.8500 | high | multi | 28 | 8 | 0 | 20 | high | under bark | -1.2467 |
| Ips sexdentatus | 0.1078 | 0.1526 | 0.0039 | 0.1482 | Curculionidae | pest | temperate | unprotected | pest | xylovorous | cambiophagous | under bark | dry and warm | fresh | coniferous | small | 5.5000 | 7.5000 | 6.5000 | high | multi | 21 | 10 | 0 | 11 | high | under bark | -0.7359 |
| Ips typographus | 0.1552 | 0.1801 | 0.0032 | 0.1106 | Curculionidae | pest | boreal | unprotected | pest | xylovorous | cambiophagous | under bark | moist and cold | fresh | coniferous | small | 4.2000 | 5.5000 | 4.8500 | high | multi | 38 | 17 | 9 | 12 | high | under bark | -0.9668 |
| Monochamus galloprovincialis | 0.1373 | 0.1649 | 0.0036 | 0.1118 | Cerambycidae | pest | boreal-temperate | unprotected | pest | xylovorous | cambioxylophagous | wood | dry and warm | fresh | coniferous | large | 12 | 25 | 18.5000 | high | uni | 7 | 3 | 0 | 4 | high | wood/cavities | 0.9437 |
| Neomida haemorrhoidalis | 0.1953 | 0.1958 | 0.0021 | 0.0580 | Tenebrionidae | rare | boreal-temperate | unprotected | neutral | fungivorous | mycetophagous | fungi | moist and cold | old | deciduous | small | 5.5000 | 6 | 5.7500 | moderate | uni | 40 | 18 | 15 | 7 | low/moderate | fungi | -0.8408 |
| Osmoderma barnabita | 0.2189 | 0.1701 | 0.0016 | -0.0370 | Scarabaeidae | rare | temperate | protected | neutral | xylovorous | saprophagous | tree cavities | moist and cold | old | deciduous | large | 22 | 32 | 27 | low | uni | 11 | 6 | 5 | 0 | low/moderate | wood/cavities | 2.1334 |
| Phaenops cyanea | 0.2229 | 0.2111 | 0.0026 | 0.0411 | Buprestidae | pest | temperate | unprotected | pest | xylovorous | cambiophagous | under bark | dry and warm | fresh | coniferous | medium | 7 | 12 | 9.5000 | high | uni | 26 | 12 | 0 | 14 | high | under bark | -0.3160 |
| Pyrochroa coccinea | 0.2605 | 0.2291 | 0.0021 | 0.0034 | Pyrochroidae | common | temperate | unprotected | neutral | invertivorous | predator & saprophagous | under bark | moist and cold | old | deciduous | large | 14 | 18 | 16 | moderate | uni | 8 | 3 | 3 | 2 | low/moderate | under bark | 0.5938 |
| Pytho depressus | 0.1267 | 0.1322 | 0.0024 | 0.0541 | Pythidae | common | boreal-temperate | unprotected | neutral | invertivorous | predator & saprophagous | under bark | moist and cold | old | various | medium | 7.5000 | 16 | 11.7500 | moderate | uni | 8 | 4 | 1 | 3 | low/moderate | under bark | -0.0011 |
| Rhagium inquisitor | 0.1285 | 0.1683 | 0.0043 | 0.1387 | Cerambycidae | common | boreal-temperate | unprotected | neutral | xylovorous | cambiophagous | under bark | moist and cold | old | coniferous | large | 10 | 21 | 15.5000 | moderate | uni | 34 | 15 | 9 | 10 | low/moderate | under bark | 0.5238 |
| Rhysodes sulcatus | 0.5153 | 0.3044 | 0.0021 | -0.3051 | Carabidae | rare | temperate | protected | neutral | xylovorous | saprophagous | wood | moist and cold | old | various | small | 6.5000 | 7.5000 | 7 | low | uni | 5 | 5 | 0 | 0 | low/moderate | wood/cavities | -0.6659 |
| Saperda scalaris | 0.2089 | 0.2245 | 0.0036 | 0.0959 | Cerambycidae | pest | temperate | unprotected | pest | xylovorous | cambioxylophagous | under bark | moist and cold | fresh | deciduous | medium | 12 | 18 | 15 | moderate | uni | 4 | 2 | 0 | 2 | low/moderate | under bark | 0.4538 |
| Schizotus pectinicornis | 0.1776 | 0.1758 | 0.0022 | 0.0462 | Pyrochroidae | common | temperate | unprotected | neutral | invertivorous | predator & saprophagous | under bark | moist and cold | old | deciduous | medium | 8 | 9 | 8.5000 | moderate | uni | 37 | 15 | 11 | 11 | low/moderate | under bark | -0.4560 |
| Scolytus intricatus | 0.2165 | 0.2257 | 0.0027 | 0.0871 | Curculionidae | pest | temperate | unprotected | pest | xylovorous | cambiophagous | under bark | dry and warm | fresh | deciduous | small | 2.5000 | 3.5000 | 3 | high | multi | 4 | 2 | 1 | 1 | high | under bark | -1.2257 |
| Spondylis buprestoides | 0.1982 | 0.2005 | 0.0029 | 0.0634 | Cerambycidae | common | temperate | unprotected | neutral | xylovorous | cambioxylophagous | wood | dry and warm | old | coniferous | large | 12 | 24 | 18 | high | uni | 7 | 3 | 0 | 4 | high | wood/cavities | 0.8737 |
| Stenurella melanura | 0.0892 | 0.1392 | 0.0061 | 0.1563 | Cerambycidae | common | temperate | unprotected | neutral | xylovorous | xylophagous | wood | moist and cold | old | various | small | 6 | 9 | 7.5000 | high | uni | 43 | 15 | 13 | 15 | high | wood/cavities | -0.5959 |
| Thanasimus formicarius | 0.1521 | 0.1885 | 0.0043 | 0.1368 | Cleridae | common | boreal-temperate | unprotected | neutral | invertivorous | predator | under bark | dry and warm | fresh | coniferous | medium | 7 | 10 | 8.5000 | high | multi | 37 | 15 | 7 | 15 | high | under bark | -0.4560 |
| Uleiota planata | 0.1498 | 0.1863 | 0.0042 | 0.1366 | Silvanidae | common | temperate | unprotected | neutral | invertivorous | predator & saprophagous | under bark | moist and cold | old | various | small | 4.5000 | 5.5000 | 5 | moderate | uni | 4 | 1 | 1 | 2 | low/moderate | under bark | -0.9458 |

**Table S3. Trait levels, species counts and eligibility for comparative testing.**

| **Trait** | **N levels** | **Min level n** | **Levels** | **Retained** |
| --- | --- | --- | --- | --- |
| Occurrence | 3 | 7 | common=10; pest=9; rare=7 | TRUE |
| Distribution | 3 | 3 | boreal=3; boreal-temperate=5; temperate=18 | FALSE |
| Conservation | 2 | 6 | protected=6; unprotected=20 | TRUE |
| Economic | 2 | 9 | neutral=17; pest=9 | TRUE |
| Diet | 3 | 3 | fungivorous=3; invertivorous=9; xylovorous=14 | FALSE |
| Shade tolerance | 2 | 8 | dry and warm=8; moist and cold=18 | TRUE |
| Settlement | 2 | 9 | fresh=9; old=17 | TRUE |
| Tree type | 3 | 7 | coniferous=10; deciduous=9; various=7 | TRUE |
| Body size | 3 | 7 | large=7; medium=9; small=10 | TRUE |
| Voltinism | 2 | 5 | multi=5; uni=21 | TRUE |
| Dispersal 2 | 2 | 11 | high=11; low/moderate=15 | TRUE |
| Microhabitat 3 | 3 | 3 | fungi=3; under bark=15; wood/cavities=8 | FALSE |
| Trophic guild | 7 | 2 | saprophagous=2 (Rhysodes, Osmoderma) etc. | FALSE |
| Host tree | 9 | 1 | 9 levels, several n<3 | FALSE |

**Table S4. Species means calculated using all-position and retained-variant-site denominators. Variant-site π is reported on a different denominator and is therefore not directly comparable in magnitude with all-position π.**

| **Species** | **Obs Het allpos** | **Exp Het allpos** | **Pi allpos** | **Fis allpos** | **Obs Het variant** | **Exp Het variant** | **Pi variant** | **Fis variant** |
| --- | --- | --- | --- | --- | --- | --- | --- | --- |
| Bolitophagus reticulatus | 0.0020 | 0.0018 | 0.0020 | 0.0001 | 0.2267 | 0.2040 | 0.2305 | 0.0118 |
| Boros schneideri | 0.0019 | 0.0019 | 0.0022 | 0.0007 | 0.1657 | 0.1711 | 0.1937 | 0.0633 |
| Chalcophora mariana | 0.0024 | 0.0025 | 0.0029 | 0.0010 | 0.2010 | 0.2113 | 0.2389 | 0.0853 |
| Cucujus cinnaberinus | 0.0022 | 0.0020 | 0.0022 | 0.0002 | 0.2187 | 0.2012 | 0.2274 | 0.0211 |
| Cucujus haematodes | 0.0025 | 0.0027 | 0.0031 | 0.0013 | 0.2074 | 0.2274 | 0.2565 | 0.1076 |
| Diaperis boleti | 0.0025 | 0.0029 | 0.0032 | 0.0016 | 0.1977 | 0.2278 | 0.2558 | 0.1290 |
| Elater ferrugineus | 0.0028 | 0.0028 | 0.0031 | 0.0006 | 0.2970 | 0.2869 | 0.3236 | 0.0625 |
| Elateroides dermestoides | 0.0022 | 0.0026 | 0.0029 | 0.0014 | 0.1792 | 0.2055 | 0.2310 | 0.1104 |
| Ips acuminatus | 0.0016 | 0.0022 | 0.0025 | 0.0021 | 0.1134 | 0.1549 | 0.1756 | 0.1446 |
| Ips sexdentatus | 0.0024 | 0.0035 | 0.0039 | 0.0034 | 0.1078 | 0.1526 | 0.1726 | 0.1482 |
| Ips typographus | 0.0024 | 0.0028 | 0.0032 | 0.0017 | 0.1552 | 0.1801 | 0.2041 | 0.1106 |
| Monochamus galloprovincialis | 0.0027 | 0.0032 | 0.0036 | 0.0022 | 0.1373 | 0.1649 | 0.1863 | 0.1118 |
| Neomida haemorrhoidalis | 0.0019 | 0.0019 | 0.0021 | 0.0006 | 0.1953 | 0.1958 | 0.2208 | 0.0580 |
| Osmoderma barnabita | 0.0018 | 0.0014 | 0.0016 | -0.0003 | 0.2189 | 0.1701 | 0.1943 | -0.0370 |
| Phaenops cyanea | 0.0025 | 0.0023 | 0.0026 | 0.0005 | 0.2229 | 0.2111 | 0.2390 | 0.0411 |
| Pyrochroa coccinea | 0.0021 | 0.0018 | 0.0021 | 0.0000 | 0.2605 | 0.2291 | 0.2600 | 0.0034 |
| Pytho depressus | 0.0020 | 0.0021 | 0.0024 | 0.0009 | 0.1267 | 0.1322 | 0.1501 | 0.0541 |
| Rhagium inquisitor | 0.0029 | 0.0038 | 0.0043 | 0.0031 | 0.1285 | 0.1683 | 0.1904 | 0.1387 |
| Rhysodes sulcatus | 0.0031 | 0.0019 | 0.0021 | -0.0019 | 0.5153 | 0.3044 | 0.3427 | -0.3051 |
| Saperda scalaris | 0.0030 | 0.0032 | 0.0036 | 0.0014 | 0.2089 | 0.2245 | 0.2522 | 0.0959 |
| Schizotus pectinicornis | 0.0019 | 0.0019 | 0.0022 | 0.0005 | 0.1775 | 0.1758 | 0.1987 | 0.0462 |
| Scolytus intricatus | 0.0023 | 0.0024 | 0.0027 | 0.0009 | 0.2165 | 0.2257 | 0.2559 | 0.0871 |
| Spondylis buprestoides | 0.0025 | 0.0026 | 0.0029 | 0.0008 | 0.1982 | 0.2005 | 0.2262 | 0.0634 |
| Stenurella melanura | 0.0035 | 0.0054 | 0.0061 | 0.0060 | 0.0892 | 0.1392 | 0.1570 | 0.1563 |
| Thanasimus formicarius | 0.0031 | 0.0038 | 0.0043 | 0.0028 | 0.1521 | 0.1885 | 0.2123 | 0.1368 |
| Uleiota planata | 0.0030 | 0.0037 | 0.0042 | 0.0027 | 0.1498 | 0.1863 | 0.2103 | 0.1366 |

**Table S5. Complete phylogenetic-signal results (Pagel’s λ and Blomberg’s K) for raw and log-transformed metrics in the full 26-species dataset and the 25-species sensitivity dataset excluding Rhysodes sulcatus.**

| **Metric** | **Scale** | **Dataset** | **N tips** | **lambda** | **lambda p** | **K** | **K p** |
| --- | --- | --- | --- | --- | --- | --- | --- |
| Fis | raw | full | 26 | 0.9379 | 0.0000 | 1.0200 | 0.0020 |
| Fis | raw | without Rhysodes sulcatus | 25 | 0.4828 | 0.0469 | 0.2717 | 0.0130 |
| He | log | full | 26 | 0.3060 | 0.4457 | 0.2683 | 0.0150 |
| He | log | without Rhysodes sulcatus | 25 | 0.0001 | 1 | 0.1871 | 0.1170 |
| He | raw | full | 26 | 0.4706 | 0.2175 | 0.3068 | 0.0130 |
| He | raw | without Rhysodes sulcatus | 25 | 0.0653 | 0.8723 | 0.1848 | 0.1230 |
| Ho | log | full | 26 | 0.6570 | 0.0296 | 0.4639 | 0.0030 |
| Ho | log | without Rhysodes sulcatus | 25 | 0.2566 | 0.2320 | 0.2410 | 0.0240 |
| Ho | raw | full | 26 | 0.9528 | 0.0004 | 0.8217 | 0.0010 |
| Ho | raw | without Rhysodes sulcatus | 25 | 0.2663 | 0.2099 | 0.2378 | 0.0250 |
| Pi | log | full | 26 | 0.5832 | 0.0099 | 0.3003 | 0.0090 |
| Pi | log | without Rhysodes sulcatus | 25 | 0.5799 | 0.0126 | 0.2857 | 0.0130 |
| Pi | raw | full | 26 | 0.4715 | 0.0318 | 0.2513 | 0.0370 |
| Pi | raw | without Rhysodes sulcatus | 25 | 0.4680 | 0.0372 | 0.2446 | 0.0450 |
